# A shared transcriptional network in the nucleus accumbens supports resilience to chronic stress across sex

**DOI:** 10.64898/2026.02.17.706464

**Authors:** Trevonn M. Gyles, Leanne M. Holt, Lyonna F. Parise, Rachel L. Fisher-Foye, Arthur Godino, Caleb J. Browne, Brian T. Kipp, Angelica M. Minier-Toribio, Tamara Markovic, Teagan Daly, Romain Durand-de Cuttoli, Long Li, Aarthi Ramakrishnan, Molly Estill, Matthew Rivera, Yun Young Yim, Astrid Carona-Acosta, Omar Sial, Chinonso A. Nwakama, Carlos A. Bolaños-Guzmán, Li Shen, Bin Zhang, Scott J. Russo, Eric M. Parise, Eric J. Nestler

## Abstract

Although chronic stress increases the risk for depression, only a subset of exposed individuals develop psychiatric illness. The biological mechanisms that protect against depression remain incompletely understood, particularly at the molecular level. Here, we identify a transcriptional network in the nucleus accumbens (NAc), a central brain reward region, that supports stress resilience in both sexes and demonstrate the causal contribution of key hub genes. Using chronic social defeat stress, RNA-seq, and co-expression network analysis, we find sex-specific but overlapping gene modules linked to resilience, anchored by shared hub genes embedded within a common network architecture. Overexpression of these hub genes in stress-naive mice confers stress protection and induces a transcriptional state that is discrete from both susceptible and resilient profiles. These findings position resilience as a structured and targetable molecular phenotype and provide a basis for investigating sex-informed mechanisms of stress adaptation.

## Introduction

Chronic stress is a major risk factor for depression, yet individuals differ widely in their responses to stress. While extensive work has characterized molecular and circuit mechanisms associated with stress susceptibility, far less is known about the biological processes that support resilience, the ability to maintain adaptive behavior despite adversity^1–5^. Recent work has expanded the ethological validity of rodent stress models, showing that social threat processing and adaptive defensive strategies are encoded by specialized neural circuits that modulate vulnerability and resilience to stress^5,6^. Understanding these mechanisms is essential for identifying targets that promote positive stress outcomes.

Sex is a critical but underexplored dimension of stress adaptation. Women are diagnosed with major depressive disorder at roughly twice the rate of men^7,8^. Recent transcriptomic studies reveal pronounced sex differences in the molecular signatures of depression in humans^9–12^. These differences are grounded in genetic, hormonal, and epigenetic mechanisms that shape sex-specific neuroendocrine and molecular adaptations across development, with recent work demonstrating that stress engages distinct regulatory pathways in males and females^13,14^. Despite this literature, we do not know whether the transcriptional architecture that supports resilience, rather than susceptibility, is similarly sex divergent or whether shared molecular programs also contribute to adaptive outcomes across sexes.

Clinical studies focused on the neurobiological mechanisms of resilience in humans is limited by the absence of standardized criteria for identifying resilient individuals^1,15^. Animal models therefore provide a powerful framework for dissecting the biological substrates that differentiate resilient from susceptible phenotypes^3–5^. Yet preclinical research on depression has mostly focused on male animals, leaving female-specific and cross-sex mechanisms of resilience insufficiently characterized.

Similarly, most preclinical studies of chronic stress have focused on the mechanisms that drive stress susceptibility, leaving stress resilience comparatively understudied^3,4,16,17^. As a result, we know less about whether the molecular substrates of resilience resemble or diverge from those that govern susceptibility, and know virtually nothing about whether resilient males and females recruit similar or distinct transcriptional programs. This knowledge is especially important given the marked sex-specific transcriptional signatures of human depression mentioned above.

In this study, we address this gap by defining the transcriptional organization of stress resilience in both male and female mice. Using the chronic social defeat stress (CSDS) paradigm adapted for use in female mice^18^, we combine RNA-sequencing with co-expression network analysis to identify gene modules across several limbic brain regions associated with resilience. This framework allows us to determine whether resilience is supported by sex-specific transcriptional pathways, sex-shared network architecture, or both. We further examine the functional relevance of key hub genes embedded within these networks to test whether they influence the behavioral and molecular features of resilience. Together, these studies provide a mechanistic foundation for understanding resilience across sexes and establish a framework for identifying regulatory pathways that promote adaptive responses to chronic stress.

## Results

### Behavioral and transcriptional stratification following CSDS

To study stress responses in females, we used a CSDS paradigm adapted for female mice (Fig. 1a)^18^. Following 10 days of CSDS, mice were classified as susceptible (SUS) or resilient (RES) using a social interaction (SI) test, which is known to correlate strongly with several other behavioral outcome measures^5,16^ Total distance traveled did not differ across groups, indicating that behavioral classification was not confounded by locomotion (Fig. 1b). SUS mice showed robust social avoidance, spending less time in the interaction zone and more time in the corners in the presence of a target mouse. By contrast, RES mice maintained social interaction levels comparable to controls (Fig. 1c,d). These behavioral phenotypes closely mirror what has been consistently observed in male mice subjected to CSDS, where SI testing reliably distinguishes susceptible and resilient subpopulations based on social avoidance behaviors^3,4,16^.

**Figure 1.**
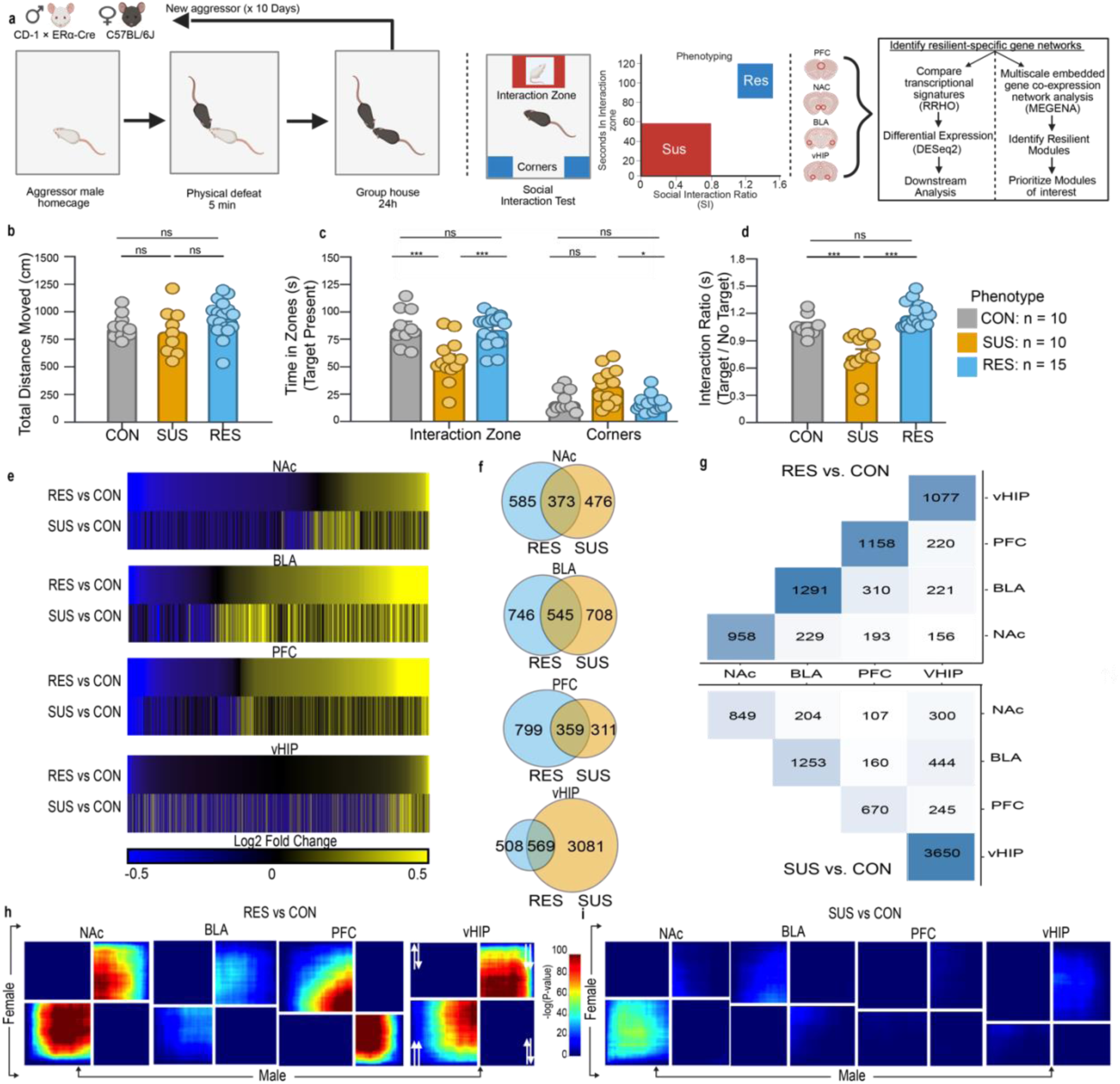
Behavioral stratification and region-specific transcriptional signatures following CSDS in female mice. a, Schematic of the adapted CSDS paradigm for female mice including the downstream RNA-seq and computational workflow: DESeq2 differential expression, MEGENA network analysis, and RRHO cross-sex comparisons. **b,** Total distance traveled during the SI test did not differ across control (CON), susceptible (SUS), and resilient (RES) groups (one-way ANOVA; ns), indicating no locomotor confound. **c,** Time spent in the interaction zone and corners during target-present trials. SUS mice spent significantly less time in the interaction zone and more time in the corners compared with CON and RES mice (one-way ANOVA with post hoc tests; ***P < 0.001; **P < 0.01; *P < 0.05). **d,** Social interaction ratio (target/no-target). SUS mice showed reduced interaction ratios relative to CON and RES mice, whereas RES mice did not differ from controls. n represents the number of mice selected for downstream sequencing stratified by phenotype. **e,** Heatmaps showing brain region–specific differential expression for RES vs. CON and SUS vs. CON comparisons across BLA, NAc, PFC, and vHIP. RES and SUS transcriptional signatures exhibit region-dependent similarity and divergence. **f,** Venn diagram quantifying the number of DEGs shared for RES vs. CON and SUS vs. CON in each brain region. **g,** Overlap matrix quantifying the number of DEGs shared across brain regions for RES vs. CON (top) and SUS vs. CON (bottom). Diagonal shading indicates total DEGs per region **h,** RRHO2 analysis comparing female and male RES vs. CON transcriptional signatures. Strong cross-sex concordance was observed in the NAc and vHIP, with minimal overlap in the PFC and BLA. **i,** RRHO2 comparison of female and male SUS vs. CON signatures revealed limited cross-sex concordance, with modest overlap restricted to the NAc. Data in **b–d** represent mean ± s.e.m.; CON: n = 10, SUS: n = 10, RES: n = 15.

To identify molecular correlates of these two behavioral phenotypes, we performed RNA-seq on four brain regions: the nucleus accumbens (NAc), prefrontal cortex (PFC), basolateral amygdala (BLA), and ventral hippocampus (vHIP) from control, susceptible, and resilient female mice. Most of the differentially expressed genes (DEGs) were unique to a single brain region, underscoring the idea that resilience and susceptibility are encoded through distinct mechanisms that vary by brain area (Fig. 1e,f,g). This finding is significant as it highlights that stress does not generate a uniform impact on the brain; rather, it induces unique transcriptional programs within several interacting circuits. Similar region-specific transcriptional distinctions between susceptible and resilient phenotypes have been observed previously in male mice exposed to CSDS, supporting the idea, along with our present findings, that behavioral divergence is accompanied by distinct molecular signatures in both sexes^3,4,16^.

To determine whether these female transcriptional signatures generalize across sex, we compared our dataset with two previously published RNA-seq studies of male CSDS^3,4^. Using rank–rank hypergeometric overlap (RRHO2)^19^, we observed cross-sex concordance in RES mice that was localized to the NAc and vHIP, while the PFC contained strikingly discordant signatures with the BLA displaying minimal similarity (Fig. 1h). In contrast, SUS transcriptional signatures showed very limited cross-sex overlap, with modest similarity restricted to the NAc and little to no shared organization in other regions (Fig. 1i). These findings suggest that resilience engages more conserved transcriptional programs across sexes compared to susceptibility which involves more sex-divergent molecular responses.

### Resilience is associated with shared transcriptional patterns in the NAc

To better understand transcriptional changes in response to chronic stress, we performed a pattern analysis on DEGs based on their expression trajectories across all samples, independent of behavioral group. This analysis, conducted separately for males and females, revealed highly similar expression patterns in the NAc (Fig. 2a,b), whereas the PFC, BLA, and vHIP showed more sex-divergent profiles (Supplemental Fig. 1). In the NAc in both sexes, four expression patterns emerged: two resilience-specific patterns (Patterns A and B), in which gene expression was selectively altered in resilient but not susceptible mice, and two stress-responsive patterns (Patterns C and D), in which gene expression changed in both resilient and susceptible mice relative to controls.

**Figure 2.**
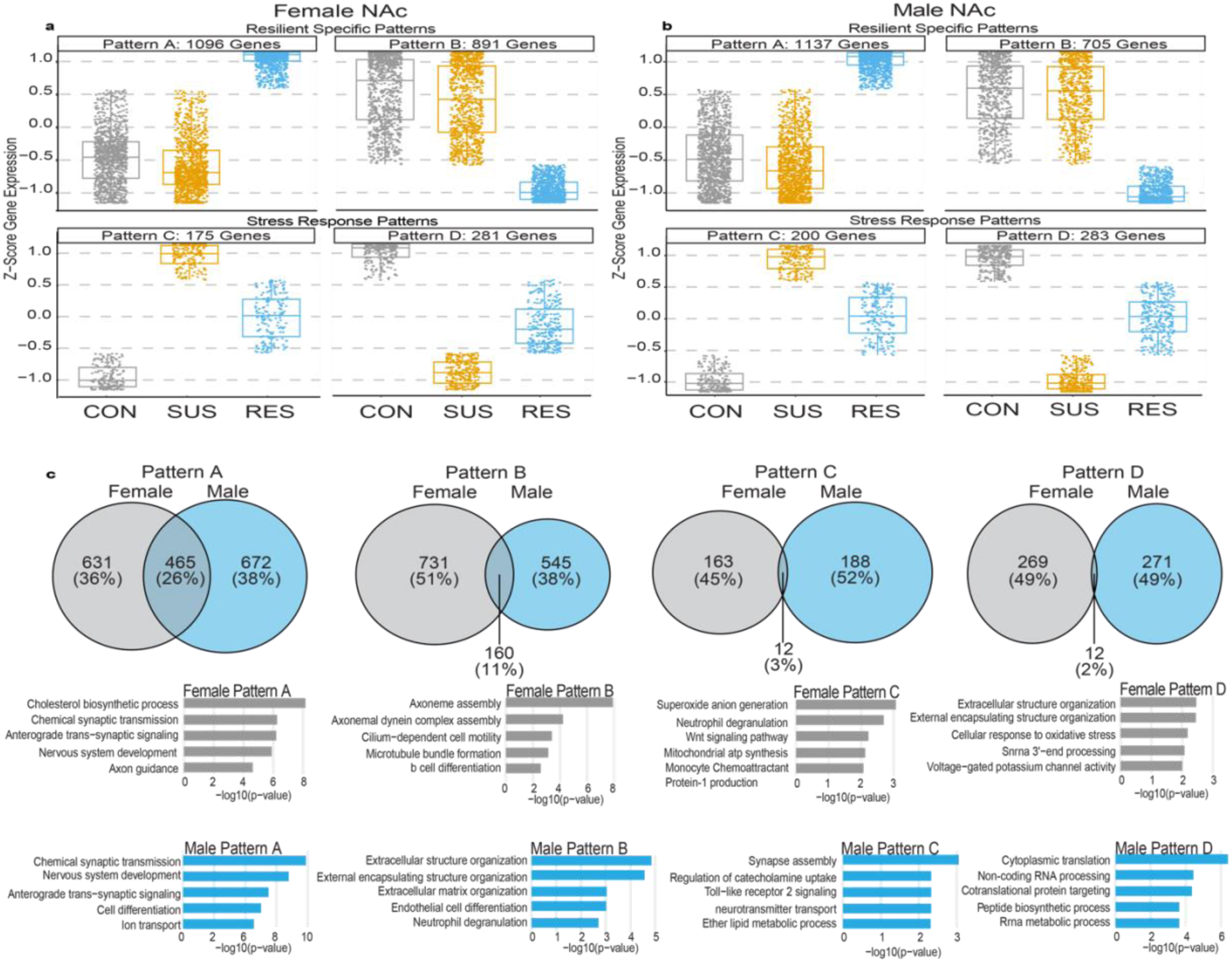
Stress resilience is associated with shared transcriptional patterns in the NAc. **a**, Pattern analysis of differentially expressed genes (DEGs) in the female NAc, showing four reproducible expression trajectories across control (CON), susceptible (SUS), and resilient (RES) groups. Patterns A and B represent resilience-specific expression changes, whereas Patterns C and D reflect stress-responsive changes shared by SUS and RES mice but with different magnitudes in the two groups. Boxplots display z-scored gene expression for each pattern. **b**, Equivalent pattern analysis performed on published male NAc CSDS datasets, revealing four analogous patterns. As in females, Patterns A and B correspond to resilience-specific changes, while Patterns C and D reflect general stress responses. **c**, Cross-sex overlap of pattern-specific gene sets. Venn diagrams show the proportion of female and male genes shared within each pattern. Resilience-specific Patterns A and B exhibit the highest cross-sex overlap (26% and 11%, respectively), whereas stress-responsive Patterns C and D show minimal overlap (<5%). Beneath each diagram, representative GO terms associated with each pattern are shown, highlighting synaptic and developmental pathways in resilience - specific patterns and immune, oxidative stress, or translational pathways in stress-responsive patterns.

Patterns A and B highlight that resilience represents a transcriptionally distinct state rather than the simple opposite of susceptibility. By contrast, Patterns C and D illustrate that resilient and susceptible mice also share a subset of gene expression changes which differ in magnitude in the two phenotypes, consistent with partially overlapping stress-responsive pathways. Cross-sex comparisons showed the greatest overlap within the resilience-specific patterns: 26% of female Pattern A genes overlapped with male Pattern A, and 11% overlapped in Pattern B (Fig. 2c). In contrast, Patterns C and D showed minimal cross-sex overlap (<5%), indicating that general stress responses are more sex-divergent than resilience-specific pathways.

Gene ontology (GO) analyses supported this distinction. Pattern A genes in both sexes were enriched for processes related to synaptic signaling, axon guidance, and neuronal development, suggesting engagement of adaptive plasticity mechanisms in resilience. Pattern B showed enrichment for cytoskeletal, microtubule, and extracellular matrix organization pathways, indicating selective downregulation of structural remodeling processes in resilient animals. Stress-responsive patterns exhibited more heterogeneous and sex-biased enrichments, including immune- and oxidative stress–related processes in females (Pattern C) and translational and metabolic pathways in males (Pattern D).

Together, these data identify the NAc as a region, unlike several others, in which resilience is defined by partially conserved transcriptional programs across sexes. The selective cross-sex overlap within resilience-specific patterns, combined with their functionally convergent GO enrichments, suggest that resilience engages shared biological pathways alongside sex-biased adaptations. These observations motivated our focus on the NAc to identify conserved regulatory mechanisms that promote adaptive stress responses.

### Co-expression network analysis reveals phenotype-specific module enrichment

To characterize how transcriptional changes are organized within the NAc, we constructed multiscale co-expression networks using MEGENA^20^ or females and males separately. We then tested each module for enrichment of DEGs from the RES vs CON and SUS vs CON comparisons and visualized significant modules using sunburst plots (Fig. 3a,b). Equivalent analyses for the PFC, BLA, and vHIP are shown in Supplemental Fig. 2.

**Figure 3.**
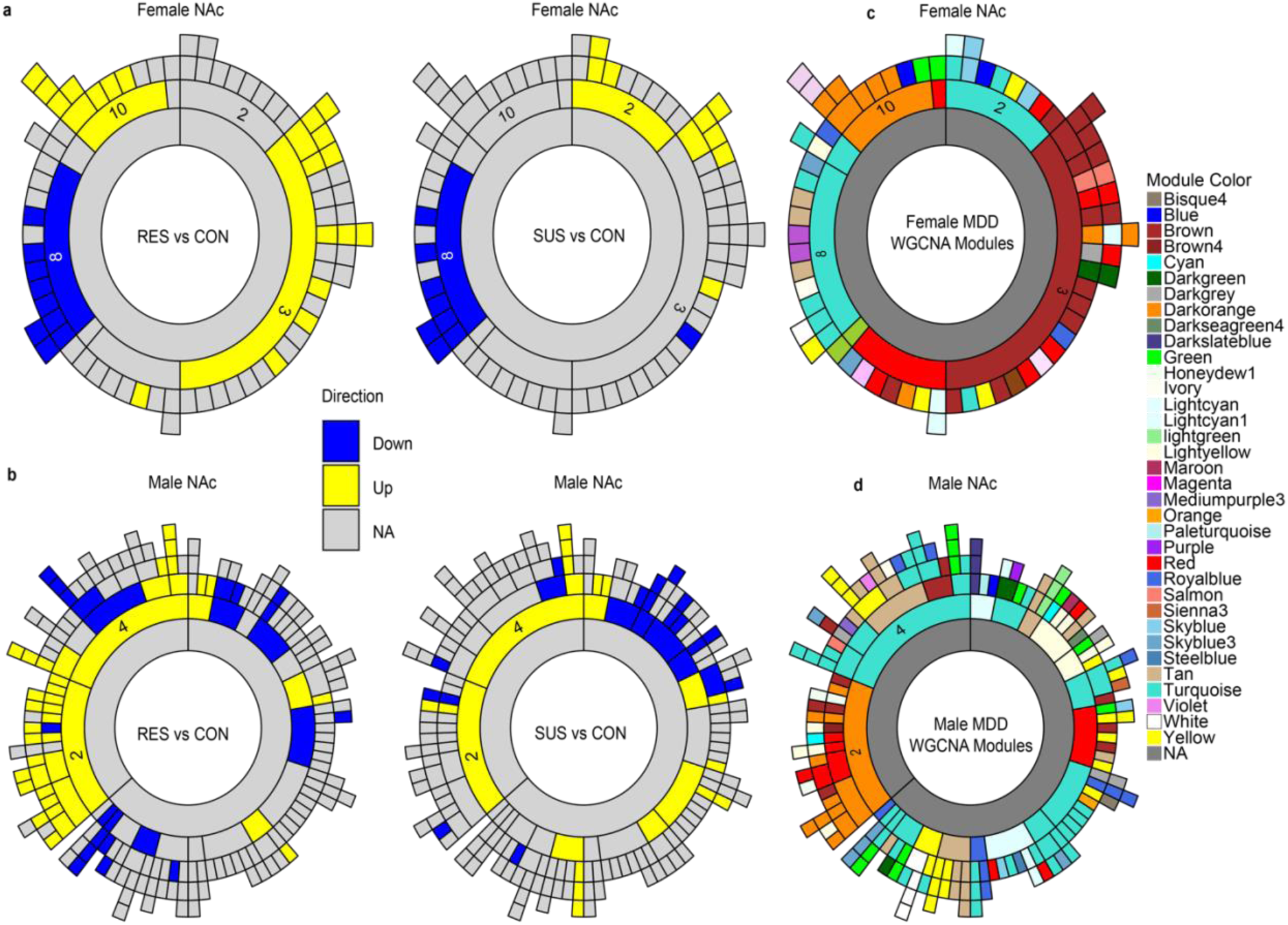
MEGENA co-expression network organization and DEG-enriched modules in the NAc of female and male mice after CSDS. **a**, MEGENA sunburst plots for the female NAc showing module enrichment for RES vs CON (left) and SUS vs CON (right). Inner rings represent parent modules and outer rings represent child modules. Upregulated DEGs in resilient mice map to child modules within parent modules 10 and 3, while susceptibility-associated upregulated DEGs map primarily to parent module 2. Downregulated DEGs in both phenotypes localize to the same parent module, 8. **b**, MEGENA sunburst plots for the male NAc for RES vs CON (left) and SUS vs CON (right). Upregulated and downregulated DEGs from both phenotypes map to similar parent modules, but resilience-associated DEGs show broader enrichment across multiple child modules compared to susceptibility-associated DEGs. **c**, Overlay of human WGCNA modules from previously published studies onto mouse MEGENA NAc networks for females (top) and males (bottom). Human-derived modules map discretely onto subsets of the mouse co-expression modules, demonstrating cross-species correspondence in network organization.

In the female NAc, resilience-associated upregulated DEGs were enriched in “child modules” belonging to “parent modules” 10 and 3 (numbers are arbitrary), whereas susceptibility-associated upregulated DEGs localized primarily to parent module 2. Downregulated DEGs from both resilient and susceptible mice mapped to the same parent module, 8. In the male NAc, RES-and SUS-associated DEGs mapped to overlapping parent modules, but resilience-associated DEGs showed broader enrichment across multiple child modules compared to susceptibility-associated DEGs. The greater number of smaller child modules observed in the male data compared to females likely reflects differences in dataset composition, including the lower number of biological replicates in the previously generated male RNA-seq datasets.

To relate mouse NAc co-expression architecture to human transcriptional networks, we overlaid previously published human WGCNA modules^9,10^ onto our MEGENA module structures (Fig. 3c). Human-derived modules mapped discretely onto subsets of mouse MEGENA modules in both sexes, and module preservation analysis revealed structured relationships within and across species (Supplemental Fig. 3). Using a preservation threshold (Zsummary ≥ 10), 24 of 187 male NAc modules (12.8%) were strongly preserved in females, whereas 5 of 85 female modules (5.8%) were preserved in males. However, these preserved module sets encompassed comparable numbers of genes: the preserved male modules together contained 6,734 genes, while the preserved female modules contained 7,245 genes. Cross-species comparisons showed a similar pattern, with 93 male modules (49.7%) and 8 female modules (9.4%) preserved in human MDD networks, corresponding to 13,183 and 6,811 mouse genes, respectively. Thus, although preservation at the module level differs between sexes, preservation at the gene level reveals substantial cross-sex and cross-species overlap in NAc co-expression architecture.

### Shared hub genes form a partially conserved resilience scaffold

To further characterize the co-expression architecture underlying resilience, we focused on the modules that showed the strongest enrichment for resilience-associated DEGs in each sex. In females, Module10 displayed the highest proportion of upregulated DEGs in the RES vs CON comparison, whereas in males Module2 showed the strongest enrichment (Supplemental Fig. 4a,b). These two clusters also showed the highest cross-sex overlap, sharing 11 percent of their gene content with significant enrichment statistics (Supplemental Fig. 4c).

Visualization of the network structures for the top 100 nodes in female Module10 and male Module2 revealed similar overall organization, with numerous shared genes occupying comparable positions within each network (Fig. 4a,b). Node coloring based on sex specificity indicated that both modules contain mixtures of female-specific, male-specific, and shared nodes. Examination of node strength distributions for the shared genes showed a broad range in both sexes, with no skew toward either high or low strength values (Supplemental Fig. 4d).

**Figure 4.**
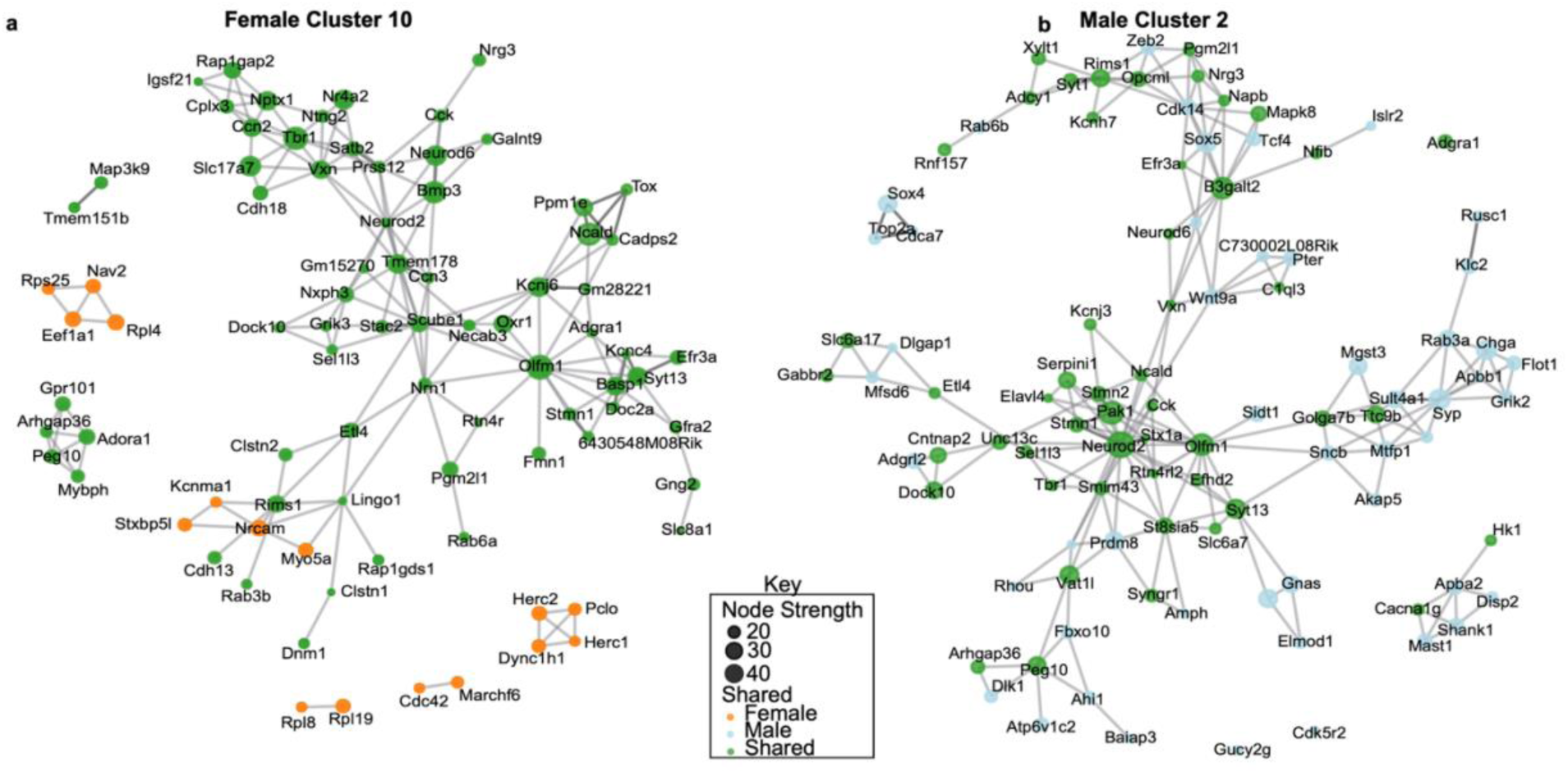
Network structure of resilience-enriched co-expression modules in NAc in female Module10 and male Module2. **a**, MEGENA subnetwork visualization of the top 100 nodes in female Module10. Edges represent co-expression connections, and node size reflects node strength. Nodes are colored according to category: female-specific (orange), male-specific (blue), or shared between sexes (green). **b**, MEGENA subnetwork visualization of the top 100 nodes in male Module2, displayed using the same node attributes and coloring scheme as in panel a. The network shows a mixture of sex-specific and shared nodes.

These results indicate that female Module10 and male Module2 contain partially overlapping sets of highly connected genes and display comparable network architecture, establishing them as the modules with the strongest cross-sex correspondence among the resilience-enriched co-expression clusters.

### Key hub genes are differentially expressed in human MDD and in stress-exposed mice

To evaluate whether the hub genes identified in the resilience-associated NAc modules show relevance across species, we first examined their expression in postmortem human NAc tissue from individuals with MDD and matched controls^9,10^. In males, *GPRIN1*, *BCR*, and *STX1A* all showed reduced expression in MDD cases compared to controls (Fig. 5a). In females, *GPRIN1* expression was decreased in MDD, whereas *BCR* and *STX1A* expression did not differ between cases and controls.

**Figure 5.**
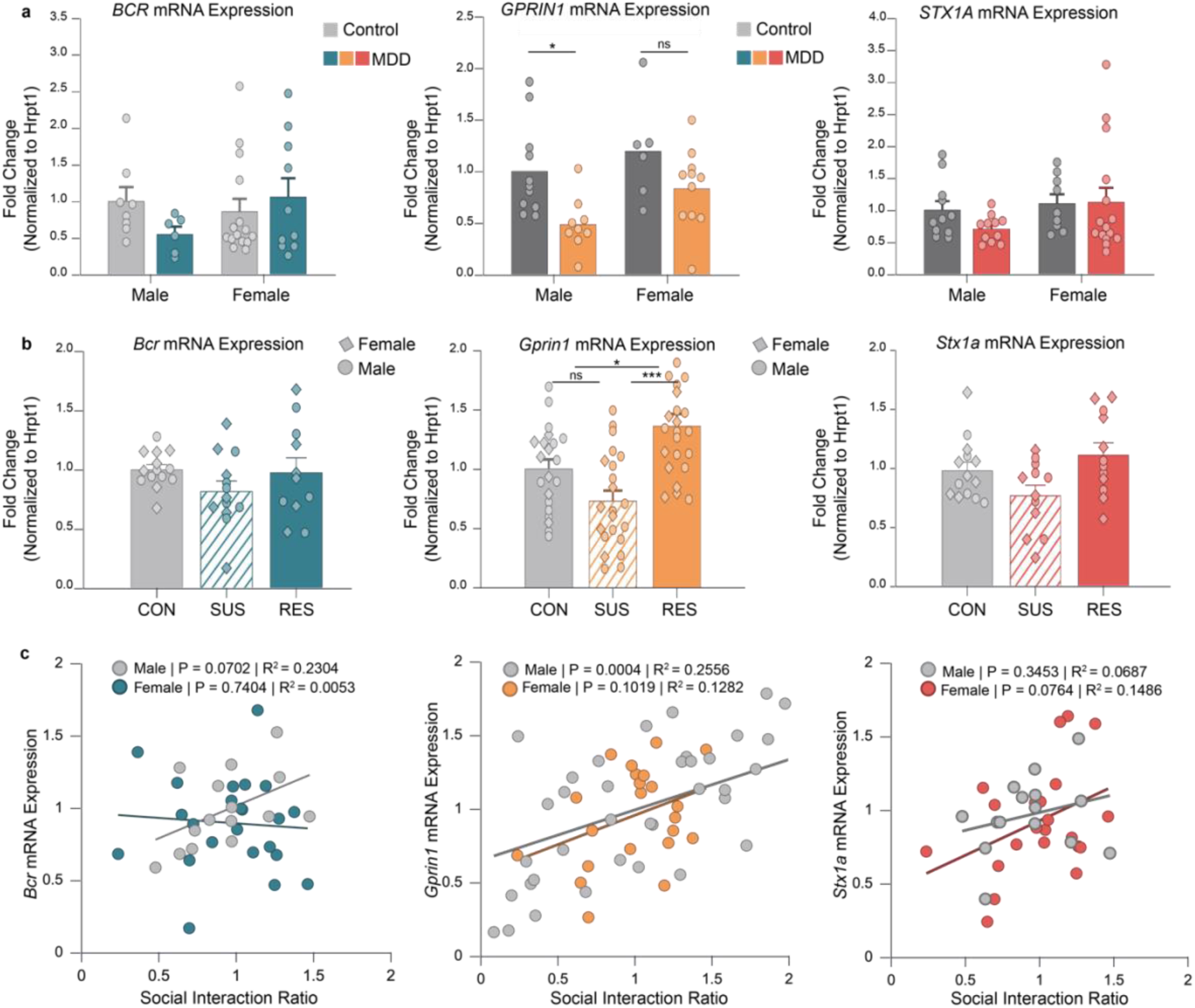
Expression of resilience-associated hub genes in the NAc of depressed humans and stress-exposed mice, and their relationship to social interaction behavior. **a**, mRNA expression of *BCR, GPRIN1*, and *STX1A* in postmortem human NAc from males and females with major depressive disorder (MDD) and matched controls. For *BCR*, a two-way ANOVA showed no significant main effects of sex (F(1,35) = 0.725, P = 0.400), diagnosis (F(1,35) = 0.341, P = 0.563), or their interaction (F(1,35) = 2.279, P = 0.140); n = 8 male controls, 6 male MDD, 15 female controls, 10 female MDD. For *GPRIN1*, a two-way ANOVA showed a significant main effect of diagnosis (F(1,33) = 10.486, P = 0.0027), with reduced expression in MDD; effects of sex (F(1,33) = 3.943, P = 0.055) and interaction (F(1,33) = 0.326, P = 0.572) were not significant; n = 11 male controls, 9 male MDD, 6 female controls, 11 female MDD. For *STX1A*, a two-way ANOVA showed no significant main effect of sex (F(1,42) = 2.129, P = 0.152), diagnosis (F(1,42) = 0.599, P = 0.443), or interaction (F(1,42) = 0.791, P = 0.379); n = 11 male controls, 11 male MDD, 9 female controls, 15 female MDD. Bars show mean ± s.e.m. **b**, mRNA expression of *Bcr, Gprin1*,and *Stx1a* in the NAc of an independent cohort of male and female mice exposed to chronic social defeat stress (CSDS). For *Bcr*, one-way ANOVA revealed no significant group differences (F(2,35) = 1.457, P = 0.247; n = 14 CON, 13 SUS, 11 RES). For *Gprin1*, one-way ANOVA revealed a significant effect of stress phenotype (F(2,63) = 12.977, P < 0.0001), with post hoc comparisons showing higher expression in RES vs CON (P = 0.016) and RES vs SUS (P < 0.0001), and no difference between CON and SUS (P = 0.090); n = 21 CON, 22 SUS, 23 RES. For *Stx1a*, a Kruskal–Wallis test indicated no significant differences among groups (H = 4.852, P = 0.088; n = 14 CON, 12 SUS, 12 RES). Bars represent mean ± s.e.m. **c**, Correlations between social interaction (SI) ratio and mRNA expression of *Gprin1*, *Bcr*, and *Stx1a* in male and female mice. For *Bcr*, regression showed no significant association in females (R² = 0.0053, P = 0.740; n = 22) and a trend in males (R² = 0.2304, P = 0.070; n = 15). For *Gprin1*, regression showed a significant positive correlation in males (R² = 0.2559, P < 0.001; n = 44) and a non-significant trend in females (R² = 0.1282, P = 0.102; n = 22). For *Stx1a*, correlations were not significant in females (R² = 0.1486, P = 0.076; n = 22) or males (R² = 0.0687, P = 0.345; n = 15). Lines represent linear fits for each sex.

We next quantified mRNA expression of these genes in the NAc of an independent cohort of male and female mice subjected to CSDS and classified as susceptible or resilient (Fig. 5b). In both sexes, *Gprin1*, *Bcr*, and *Stx1a* expression was reduced in susceptible mice relative to controls and was elevated in resilient mice. To assess the relationship between gene expression and behavioral outcomes, we examined correlations between SI ratio and mRNA levels for each gene (Fig. 5c). *Gprin1* expression showed positive correlations with SI ratio in both males and females. *Bcr* expression was positively correlated with SI ratio in males, while *Stx1a* expression was positively correlated with SI ratio in females.

### Gene overexpression produces stress-protective behavior and distinct transcriptional states in males and females

To test whether resilience-associated hub genes can actively shape behavioral responses to stress, we overexpressed *Bcr*, *Gprin1*, or *Stx1a* individually in the NAc of stress-naïve mice, allowed four weeks for expression, and then subjected animals to CSDS followed by SI testing. Since all three of these genes are highly enriched in neurons, we used an AAV vector that selectively targets neurons. In females, mCherry-expressing controls showed the expected decrease in time spent in the interaction zone and interaction ratio after CSDS, whereas mice overexpressing each of the three genes maintained normal social interaction behavior (Fig. 6a), indicating that all three genes in this brain region promote resilience. For time in the interaction zone, two-way ANOVA (virus × stress) revealed a significant main effect of virus (F(3,50) = 5.80, P = 0.0018) and no main effect of stress or interaction. Bonferroni post hoc tests showed that CSDS significantly reduced interaction time only in mCherry mice (P = 0.044) and that stressed mCherry mice spent less time in the interaction zone than *Bcr-*, *Gprin1-*, or *Stx1a*-overexpressing mice (all P < 0.001). For interaction ratio, two-way ANOVA revealed main effects of virus (F(3,51) = 5.83, P = 0.0017) and stress (F(1,51) = 8.54, P = 0.0052) but no interaction; post hoc tests again showed a decrease in mCherry mice after CSDS (P = 0.031) and higher ratios in stressed overexpression groups relative to stressed mCherry controls (all P ≤ 0.0019).

**Figure 6.**
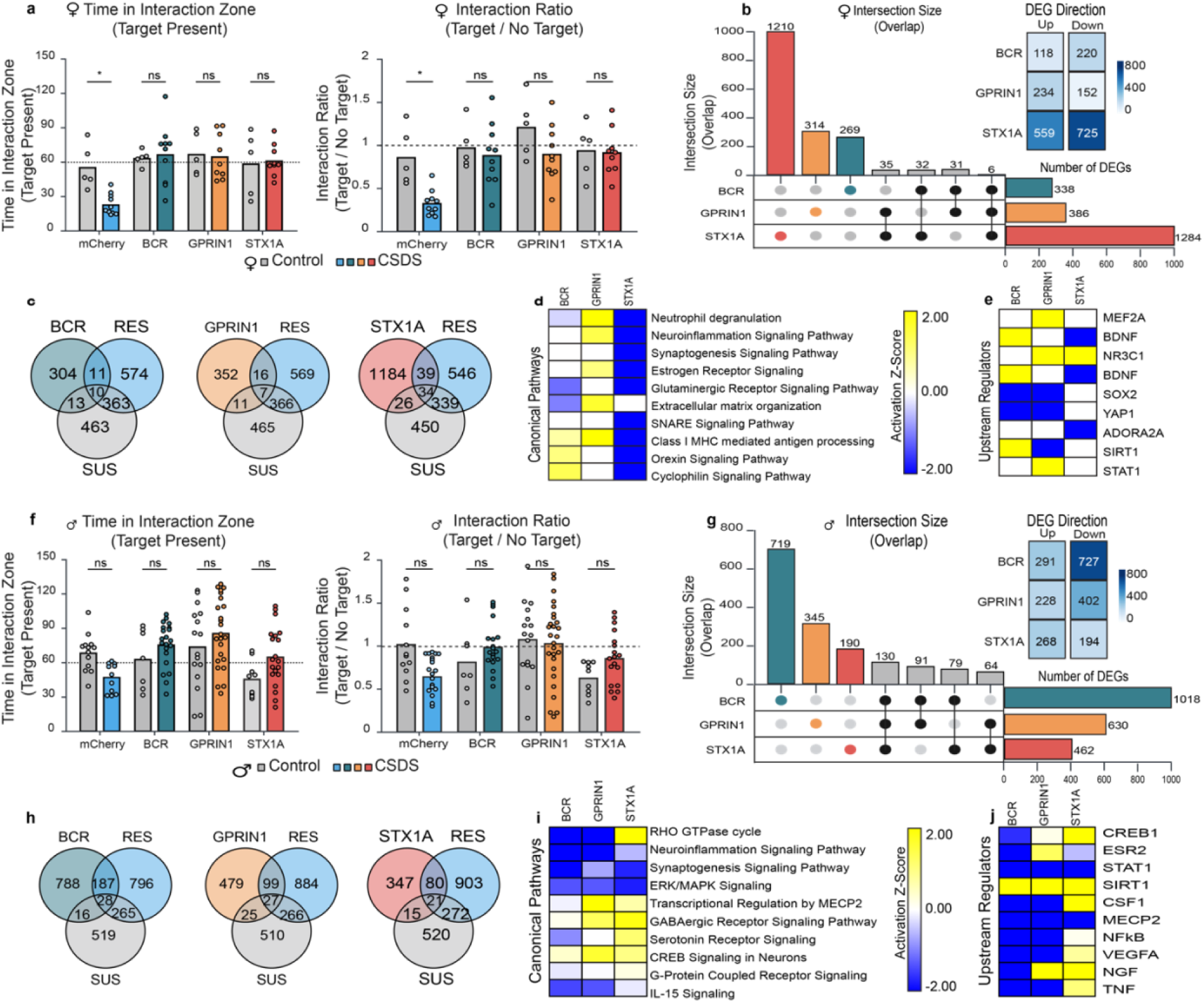
Overexpression of *Bcr*, *Gprin1*, or *Stx1a* in the NAc of female and male mice promotes behavioral resilience and distinct transcriptional profiles. **a**, Social interaction (SI) behavior in female mice following NAc overexpression of *Bcr*, *Gprin1*, or *Stx1a* compared with mCherry controls. Time spent in the interaction zone when the target is present (left) and interaction ratio (target/no target, right) are shown. Two-way ANOVAs (virus × stress) revealed significant main effects of virus on interaction time (F(3,50) = 5.8006, P = 0.0018) and interaction ratio (F(3,51) = 5.8253, P = 0.0017), and a main effect of stress on interaction ratio (F(1,51) = 8.5383, P = 0.0052). Bonferroni post hoc tests showed that CSDS reduced interaction time and ratio in mCherry mice (P = 0.044 and P = 0.0313, respectively), whereas none of the overexpression groups differed between control and CSDS conditions, and stressed mCherry mice showed lower interaction time and ratios than *Bcr-*, *Gprin1-*, or *Stx1a-*overexpressing mice (all P ≤ 0.0019). Bars show mean ± s.e.m. (n = 5 control and 10 CSDS per virus group). **b**, UpSet plot summarizing DEGs induced by each overexpression condition in female NAc. *Stx1a* produced 1,283 DEGs (559 upregulated, 725 downregulated), *Gprin1* produced 386 DEGs (234 up, 152 down), and *Bcr* produced 338 DEGs (118 up, 220 down); most DEGs are unique to each overexpressed gene. **c**, Venn diagrams showing overlap between overexpression-induced DEGs and DEGs associated with natural resilience (RES) or susceptibility (SUS) in females. **d**, Canonical pathways (IPA) enriched among DEGs for each female overexpression condition. **e**, Predicted upstream regulators (IPA) for DEGs associated with each female overexpression condition. **f**, Social interaction behavior in male mice following NAc overexpression of *Bcr*, *Gprin1*, or *Stx1a* compared with mCherry controls. Time in the interaction zone (left) and interaction ratio (right) are shown. Two-way ANOVAs (virus × stress) revealed significant main effects of virus on interaction time (F(3,113) = 6.0545, P = 0.00074) and interaction ratio (F(3,118) = 4.0961, P = 0.0083), and significant virus × stress interactions for both measures (time: F(3,113) = 3.6979, P = 0.0139; ratio: F(3,118) = 4.1381, P = 0.0079). Tukey post hoc tests showed that stressed mCherry mice had lower interaction ratios than *Gprin1*-overexpressing mice (P = 0.021) and reduced interaction times compared to *Bcr-* and *Gprin1*-overexpressing mice after CSDS (P = 0.049 and P = 0.00083, respectively), whereas overexpression groups did not differ significantly from their corresponding unstressed controls. Bars show mean ± s.e.m. (n values shown in the figure). **g**, UpSet plot of DEGs induced by overexpression in male NAc. *Stx1a* produced 998 DEGs (291 upregulated, 727 downregulated), *Gprin1* produced 630 DEGs (228 up, 402 down), and *Bcr* produced 462 DEGs (268 up, 194 down). **h**, Overlap between male overexpression-induced DEGs and natural RES or SUS DEGs. **i**, Canonical pathways (IPA) enriched among DEGs for each male overexpression condition. **j**, Predicted upstream regulators (IPA) for DEGs associated with each male overexpression condition.

We next characterized the downstream transcriptional consequences of each manipulation in the NAc using RNA-seq. In females, *Stx1a* overexpression induced the largest transcriptional response, with 1,283 DEGs (559 upregulated, 725 downregulated; Fig. 6b). *Gprin1* and *Bcr* overexpression each produced smaller but still substantial DEG sets (*Gprin1*: 386 DEGs, 234 up and 152 down; *Bcr*: 338 DEGs, 118 up and 220 down). UpSet analysis showed that most DEGs were unique to each overexpressed gene, with relatively few genes shared across manipulations (Fig. 6b). Comparison with DEGs from naturally resilient and susceptible females revealed that each overexpression condition generated a transcriptional profile that only partially overlapped with natural RES or SUS signatures (Fig. 6c); the majority of DEGs in overexpressing mice were not differentially expressed in either natural phenotype. Canonical pathway and upstream regulator analyses revealed that each gene induced a partially distinct pattern of pathway engagement in both sexes, with *Stx1a* generally producing an inhibitory tone (Fig. 6d,e).

In males, a similar pattern emerged, with overexpression of each gene promoting resilience, i.e., preventing CSDS-induced reductions in social interaction (Fig. 6f). For time in the interaction zone, two-way ANOVA (virus × stress) revealed significant main effects of virus (F(3,113) = 6.05, P = 0.00074) and a virus × stress interaction (F(3,113) = 3.70, P = 0.014). Tukey post hoc tests indicated that CSDS reduced interaction time in mCherry mice compared to their unstressed counterparts (P ≈ 0.30, trend) and that stressed mCherry mice spent less time in the interaction zone than *Bcr-* and *Gprin1-*overexpressing mice after CSDS (P = 0.049 and P = 0.00083, respectively). For interaction ratio, two-way ANOVA showed significant main effects of virus (F(3,118) = 4.10, P = 0.0083) and a virus × stress interaction (F(3,118) = 4.14, P = 0.0079). Tukey post hoc tests revealed lower interaction ratios in stressed mCherry mice compared with *Gprin1-*overexpressing mice in both stressed and unstressed conditions (P = 0.021 and 0.021, respectively), whereas overexpression groups did not differ from their respective unstressed controls.

In males, overexpression again produced gene-specific transcriptional signatures (Fig. 6g). *Stx1a* overexpression yielded 998 DEGs (291 upregulated, 727 downregulated), *Gprin1* produced 630 DEGs (228 up, 402 down), and *Bcr* produced 462 DEGs (268 up, 194 down). As in females, the DEG sets were largely distinct across genes, with limited overlap in the UpSet plot. Overlap with male natural RES and SUS DEGs was similarly modest (Fig. 6h), indicating that each overexpression condition produced a transcriptional state that was related to, but not equivalent to, naturally occurring stress phenotypes. Canonical pathway and upstream regulator analyses revealed that each gene induced a partially distinct pattern of pathway engagement in both sexes, with *Bcr* generally producing an inhibitory tone. (Fig. 6d,e).

### Overexpression-induced DEGs are largely sex-specific and map onto resilience-associated co-expression modules

To determine whether overexpression of the same hub gene in NAc affects similar transcriptional targets in males and females, we compared DEGs across sex for each viral construct. Venn diagram analyses revealed minimal overlap between female and male DEGs for all three overexpressed genes when compared to naturally resilient mice (Fig. 7a,c). Whereas 218 genes were shared between males and females in the naturally resilient condition, only 5, 50, and 28 genes were shared across sex following overexpression of *Bcr*, *Gprin1*, or *Stx1a*, respectively. Thus, overexpression of each gene elicited largely sex-specific transcriptional responses, indicating that manipulation of the same hub gene engages distinct molecular targets in males and females.

**Figure 7.**
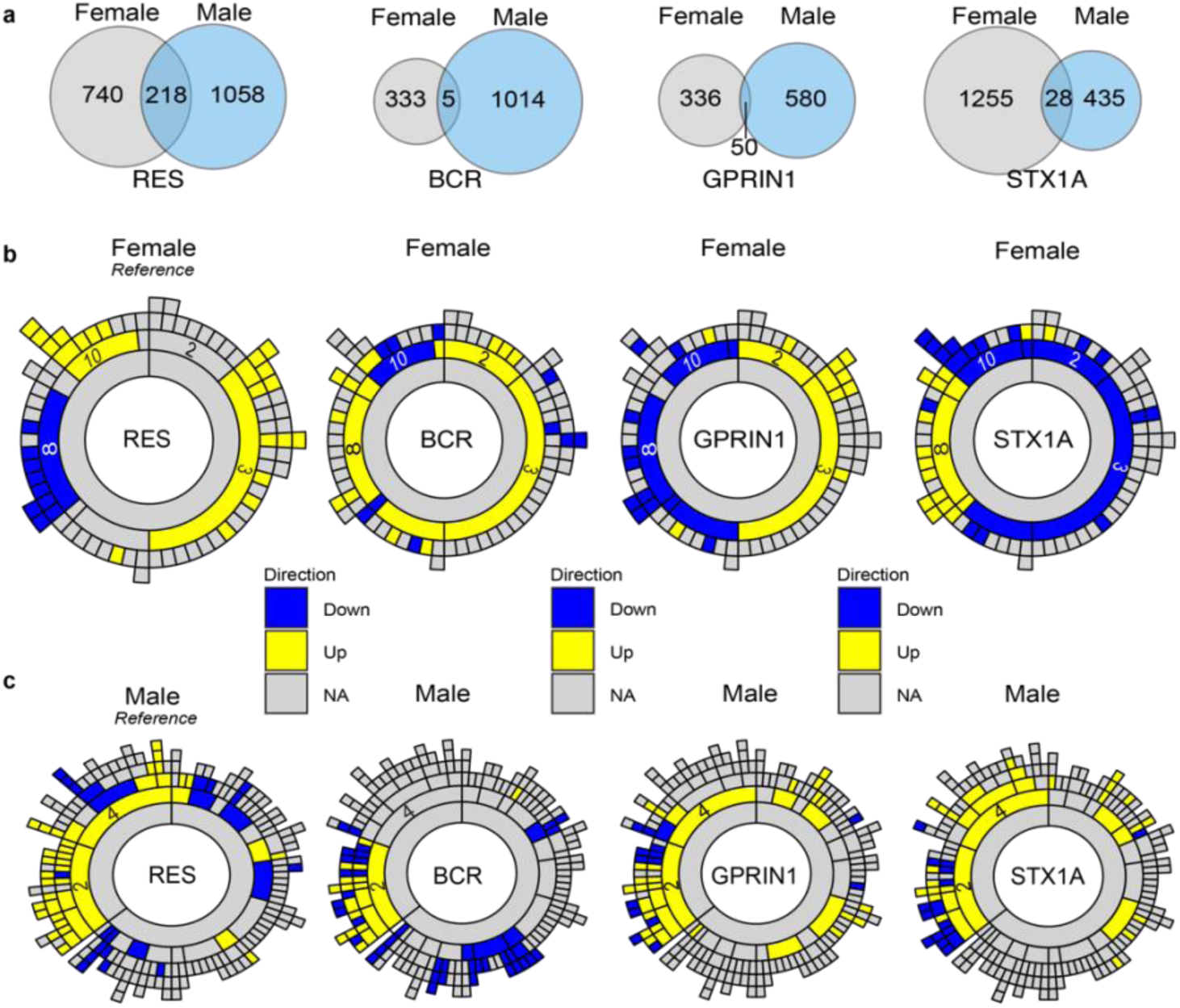
Cross-sex comparison of overexpression-induced DEGs and enrichment within resilience-associated MEGENA modules in NAc. **a**, Venn diagrams showing overlap of DEGs between female and male mice for each viral construct. For naturally resilent mice, 218 genes were shared between sexes *Bcr* = 5; *Gprin1* = 50 genes; *Stx1a* = 28 genes, indicating largely sex-specific transcriptional responses. **b**, MEGENA sunburst plots showing enrichment of female overexpression DEGs within the original NAc co-expression modules. Yellow indicates upregulated DEGs; blue indicates downregulated DEGs. **c,** MEGENA sunburst plots showing enrichment of male overexpression DEGs within the original NAc modules. Yellow indicates upregulated DEGs; blue indicates downregulated DEGs.

We next examined how overexpression-induced transcriptional changes in the NAc relate to the resilience-associated co-expression architecture defined by MEGENA. In females, overexpression of *Bcr*, *Gprin1*, or *Stx1a* produced marked shifts in the direction and distribution of module-level enrichment (Fig. 7b). *Bcr* overexpression was associated with a predominantly upregulated transcriptional profile, with enrichment of upregulated DEGs in modules 2, 3, and 8 accompanied by downregulation of module 10. *Gprin1* overexpression elicited a more balanced transcriptional response, with upregulated enrichment in modules 2 and 3, downregulated enrichment in module 8, and downregulation of module 10. In contrast, *Stx1a* overexpression produced a broadly inhibitory transcriptional profile, characterized by downregulation across modules 10, 2, and 3, together with upregulation of genes within module 8. Notably, overexpression of all three genes resulted in downregulation of module 10, the module that showed the strongest enrichment for upregulated DEGs in naturally resilient mice. Thus, although viral overexpression engages the same network modules associated with resilience, it does so by reshaping the directional balance of transcriptional activity across these modules.

In males, DEG enrichment was more tightly focused within the core resilience-associated module identified previously. Overexpression of *Bcr*, *Gprin1*, or *Stx1a* produced upregulated and downregulated DEGs that localized predominantly to parent module 2 in the male MEGENA network (Fig. 7d). The pattern of enrichment across module 2 closely resembled that observed in naturally resilient males, indicating that, although the individual DEGs are largely distinct, overexpression of these hub genes re-engages a similar network-level transcriptional signature.

Together, these findings indicate that overexpression of individual hub genes produces transcriptional responses that are largely sex-specific and distinct from naturally occurring resilient or susceptible states at the level of individual DEGs. However, despite this limited gene-level overlap, the transcriptional consequences of each overexpression condition consistently engaged co-expression modules that occur with natural resilience. Thus, while viral overexpression reshapes the direction and balance of transcriptional activity within these modules and introduces novel gene-level changes, it nevertheless recapitulates key elements of the resilience-associated network architecture identified by MEGENA. This convergence at the module level, in the context of divergent gene-level signatures, underscores the importance of network organization in shaping adaptive responses to chronic stress.

## Discussion

Stress resilience is increasingly recognized as a unique, active biological process, yet the molecular organization that supports such adaptive responses to chronic stress remains incompletely understood^5,15^. Here we leverage a sex-balanced CSDS paradigm, transcriptomic profiling across four limbic regions, co-expression network analysis, and causal gene manipulations in the NAc to identify both shared and sex-specific components of the resilience architecture. Across analyses, resilience emerges as a dynamic, structured transcriptional state not merely the absence of susceptibility.

A prominent finding from our study is that the transcriptional signature of resilience is more conserved across sex than that of susceptibility. Whereas susceptibility-associated changes in gene expression showed very little cross-sex concordance across all regions, resilience in male and female mice converged specifically within the NAc and, to a lesser degree, the vHIP. Pattern analysis further revealed that resilience-specific expression trajectories (Patterns A and B) were substantially more shared across sex than general stress-responsive trajectories (Patterns C and D). These observations suggest that resilience involves a partially conserved molecular program, while susceptibility may arise from more divergent or heterogeneous transcriptional pathways. This lack of concordance for susceptibility to CSDS in mice is consistent with similar, very little cross-sex concordance in human MDD^9–12,21^. An important goal for future work is to confirm greater cross-sex concordance for stress resilience in humans, however, this is technically challenging given the difficulty of capturing stress resilience in postmortem human brain collections.

Our network analyses using MEGENA provided a more fine-grained view of the transcriptional architecture of stress resilience. Despite differences in network size and modular structure across sex, both female and male NAc networks contained modules strongly enriched for resilience - associated DEGs. Female Module10 and male Module2 exhibited the highest enrichment and shared 11 percent of their gene content. Network visualizations revealed that these clusters also share multiple hub genes with comparable node strengths, indicating that the cross-sex overlap extends beyond individual DEGs to include similarly positioned, highly connected regulatory nodes. Although the absolute degree of overlap is modest, the preservation of connectivity patterns and the shared representation of hub genes point to a partially conserved “resilience scaffold” that is flexibly tuned by sex-specific subnetworks. The strong preservation of these male and female modules within human co-expression networks further underscores their translational relevance.

The identification and causal testing of *Stx1a*, *Gprin1*, and *Bcr* allowed us to probe how distinct nodes within the resilience-associated network contribute to adaptive stress responses. Prior work from our group identified *Stx1a* and *Gprin1* as hub genes within a transcriptional network associated with resilience in the NAc, providing an initial systems-level rationale for their selection^3,4^. Beyond this network context, *Stx1a* encodes a core component of the presynaptic SNARE complex involved in synaptic vesicle docking and neurotransmitter release, processes that have been repeatedly implicated in stress adaptation and mood-related pathology^22,23^. GPRIN1 is linked to neurite outgrowth, synaptic plasticity, and excitatory signaling, and emerging evidence suggests that it participates in stress- and activity-dependent transcriptional programs relevant to psychiatric phenotypes^21,24,25^. BCR has relatively limited prior association with mood disorders, though early candidate-gene studies and its known roles in cytoskeletal and small GTPase signaling suggest potential relevance to stress-induced network remodeling^26^.

Our cross-species analyses further highlight that the functional relevance of these genes is strongly shaped by sex. In postmortem human NAc, *GPRIN1*, *BCR*, and *STX1A* were each reduced in males with MDD, whereas only *GPRIN1* was reduced in females, suggesting sex-biased engagement of these molecular pathways in depression. This pattern was mirrored in mice, where *Gprin1* expression tracked resilience in both sexes, while *Bcr* and *Stx1a* showed sex-specific relationships with social interaction behavior, with *Bcr* more strongly associated with male outcomes and *Stx1a* with female outcomes. Despite the relatively modest strength of prior human genetic or transcriptomic associations, targeted overexpression of each gene in the NAc was sufficient to confer protection against stress-induced social avoidance, establishing a causal role for these genes in promoting resilience behavior. Notably, although each manipulation produced largely distinct and sex-specific transcriptional programs, all converged on previously identified resilience-associated network modules, indicating that sex-biased molecular implementations can nevertheless engage shared network architecture to promote adaptive behavioral outcomes.

Although overexpression of *Stx1a*, *Gprin1*, or *Bcr* in NAc was sufficient to prevent stress-induced social avoidance in both sexes, the transcriptional consequences of each manipulation were strikingly distinct. Across males and females, the downstream DEG profiles associated with each gene showed minimal overlap with one another and only partial overlap with naturally occurring resilient or susceptible states. Moreover, the magnitude and direction of these effects differed by sex, with *Stx1a* producing the largest transcriptional footprint in females and *Bcr* exerting the strongest effects in males. Notably, despite targeted overexpression, all three manipulations were accompanied by substantial downstream gene downregulation, raising the possibility that resilience-promoting perturbations may engage compensatory or homeostatic transcriptional responses rather than simple linear amplification of gene expression programs.

Comparison of overexpression-induced transcriptional signatures across sex further underscored their specificity. In contrast to the substantial cross-sex overlap observed in naturally resilient mice, overexpression of the same gene in males and females yielded largely non-overlapping DEG sets, indicating that each manipulation generates a distinct, sex-specific transcriptional state. Yet, despite this gene-level divergence, the downstream transcriptional changes consistently localized to previously identified resilience-associated co-expression modules, alongside additional effects in non-resilience modules. This pattern suggests that while overexpression creates novel transcriptional configurations that differ across genes and sexes, the core resilience signature is retained at the level of network organization. Together, these findings indicate that resilience is not encoded by a fixed set of genes, but rather by a modular network architecture that can be engaged through multiple, sex-dependent molecular routes. These results emphasize the importance of including both sexes in mechanistic studies: sex-biased implementations of a shared architecture may help reconcile why some molecular findings replicate poorly across sex while behavioral phenotypes do not.

Our study has several limitations that point to important future directions. In the present work, we overexpressed individual hub genes independently and at relatively high levels, an approach that allowed us to establish causal sufficiency but does not fully recapitulate the coordinated regulation observed within endogenous transcriptional networks. Future studies using multiplexed transcriptional modulation strategies may enable simultaneous and more physiologically relevant manipulation of multiple network components, providing a deeper understanding of how resilience-associated gene programs are orchestrated in vivo. In addition, our findings reveal clear sex-specific relationships between gene expression and behavioral outcomes, with STX1A showing stronger associations in females and BCR in males. Further investigation into how these gene-by-sex interactions influence other forms of stress exposure, social behavior, and adaptive responses will be important for defining the broader generalizability of resilience mechanisms. Finally, although our results are supported by human transcriptional data, extending these findings to human-derived cellular systems or longitudinal clinical studies will be necessary to fully establish their translational relevance.

In summary, our results support a model in which stress resilience is governed by a modular, partially conserved transcriptional architecture anchored by shared hub genes and flexibly implemented through sex-specific regulatory pathways. Distinct molecular perturbations can engage this scaffold to produce behavioral resilience, highlighting opportunities for therapeutic strategies that target network-level processes rather than individual risk genes. By integrating sex-balanced behavioral assays, computational analyses, and causal manipulations, this study provides a framework for dissecting the molecular basis of resilience and opens new avenues for developing next-generation interventions for stress-related psychiatric disorders.

## Methods

### Animals

All procedures were approved by the Institutional Animal Care and Use Committee at the Icahn School of Medicine at Mount Sinai and were conducted in accordance with NIH guidelines. Wild-type C57BL/6J mice (The Jackson Laboratory) of both sexes were used for all experiments. Unless otherwise indicated, mice were group-housed (2–5 per cage) in a temperature- and humidity-controlled vivarium on a 12 h light/dark cycle (lights on at 07:00) with food and water available ad libitum. Animals were acclimated to the facility for at least 1 week prior to any procedures. Age-matched cohorts were used within each experiment.

### Female chronic social defeat stress

Female CSDS was performed using DREADD-induced aggression in Esr1-Cre CD1 male mice, adapted from previously described protocols^18^. Briefly, Esr1-Cre CD1 males were stereotaxically injected bilaterally with AAV-DIO-hM3Dq into the ventrolateral subdivision of the ventromedial hypothalamus (VMHvl) and allowed to recover and express virus for ≥3 weeks. Aggressor males were screened for robust attack latency and maintained in individual cages. For each defeat episode, an experimental C57BL/6J female mouse was placed into the home cage of a VMHvl-hM3Dq CD1 male for 5 min of physical interaction. DREADD–mediated activation of *Esr1*+ neurons was achieved using clozapine-N-oxide (CNO; Sigma), administered intraperitoneally prior to defeat sessions, following the dosing schedule of the original report^18^. After each 5 min defeat bout, the female test mouse was housed across a perforated divider from the aggressor for the remainder of the 24 h period to allow sensory contact without further physical interaction. This procedure was repeated for 10 consecutive days with a novel aggressor each day. Control mice were housed in pairs and handled daily but were not exposed to defeat.

### Social interaction test and behavioral stratification

Social interaction (SI) testing was performed 24 h after the final defeat session in a square open field arena (44 × 44 cm) under dim red light (∼10 lux), according to standard CSDS procedures^16,17^. The arena contained a wire mesh or Plexiglas enclosure centered along one wall. During the first 2.5 min “no-target” trial, the experimental mouse freely explored the arena with an empty enclosure. During the second 2.5 min “target-present” trial, a novel, age- and sex-matched C57BL/6J mouse was placed inside the enclosure. Time spent in a predefined “interaction zone” (7.5 cm band surrounding the enclosure) and time spent in corner zones were quantified using automated video tracking (e.g. EthoVision XT; Noldus). An SI ratio was calculated as time in the interaction zone with the target present divided by time in the interaction zone with the target absent. Based on established criteria (refs), mice with SI ratio < 0.9 were classified as susceptible (SUS), mice with SI ratio > 1.1 as resilient (RES), and intermediate animals were excluded from transcriptomic analyses.

### Viral vectors and stereotaxic surgery

For gain-of-function experiments, we used AAV9 vectors encoding mCherry alone (AAV9-hSyn-mCherry; control) or mCherry-tagged mouse *Bcr*, *Gprin1*, or *Stx1a* under a neuronal promoter (AAV9-hSyn-BCR-mCherry, AAV9-hSyn-GPRIN1-mCherry, AAV9-hSyn-STX1A-mCherry; custom prepared by the Duke University Viral Vector Core in Durham, North Carolina, USA., titer ∼10^1^²–10^1^³ vg/mL).

Stereotaxic surgeries were performed as described previously^4,27^. Mice were anesthetized with ketamine (100 mg/kg, i.p.) and xylazine (10 mg/kg, i.p.) and placed in a small-animal stereotaxic frame (Kopf Instruments). Following scalp incision and craniotomy, 33-gauge Hamilton syringes were lowered bilaterally into the NAc using the following coordinates relative to Bregma: anteroposterior +1.3 mm, mediolateral ±1.5 mm, dorsoventral −4.4 mm at a 10° angle. Virus (0.5 μL per side) was infused at 0.1 μL/min. Needles were left in place for an additional 5 min to minimize backflow and then slowly withdrawn. Animals received postoperative analgesia (e.g. meloxicam) and were monitored daily. Behavioral experiments began ≥3–4 weeks after surgery to allow for stable viral expression.

### Experimental timelines for overexpression cohorts

For overexpression experiments, male and female C57BL/6J mice received bilateral NAc injections of AAV9-mCherry, AAV9-BCR-mCherry, AAV9-GPRIN1-mCherry or AAV9-STX1A-mCherry. After recovery and expression, mice were either left unstressed or subjected to 10 days of CSDS as described above. SI testing was performed 24 h after the final defeat episode. A subset of animals was perfused or decapitated 24 hours after SI for NAc tissue collection for RNA extraction and qPCR or RNA-seq.

### Human postmortem NAc RNA-seq datasets

Human NAc transcriptional data were obtained from previously published RNA-seq studies of individuals with MDD and psychiatrically healthy controls^9,10^. For the present analyses, we used the authors’ processed count matrices and module assignments (WGCNA networks) as reported. Differential expression statistics for *GPRIN1*, *BCR* and *STX1A*, as well as human co-expression modules, were used to (i) compare case–control regulation of hub genes and (ii) assess module-level overlap and preservation with mouse MEGENA networks.

### Tissue collection and NAc dissections

For mouse molecular experiments, brains were rapidly removed following cervical dislocation or deep anesthesia and decapitation. Coronal sections (1 mm) containing the NAc were cut using a brain matrix on an ice-cold surface. Bilateral NAc (core and medial shell) punches were collected using a 14-gauge tissue punch, flash frozen on dry ice, and stored at −80°C until processing. For multi-region RNA-seq (female CSDS cohort), additional punches were collected from the medial prefrontal cortex (PFC), basolateral amygdala (BLA) and ventral hippocampus (vHIP) using standard stereotaxic landmarks.

### RNA extraction and quantitative real-time PCR

Total RNA was extracted from frozen punches of the nucleus accumbens (NAc) or other brain regions using the RNeasy Micro Kit (Qiagen) according to the manufacturer’s instructions, including on-column DNase digestion. RNA concentration and purity were assessed by spectrophotometry, and samples with A260/280 ratios of approximately 2.0 were used. cDNA was synthesized from 500–1,000 ng total RNA using the iScript cDNA Synthesis Kit (Bio-Rad). Quantitative PCR (qPCR) was performed using TaqMan Gene Expression Assays (Applied Biosystems) with TaqMan Fast Advanced Master Mix on a QuantStudio 5 Real-Time PCR System (Applied Biosystems). Reactions were run in technical triplicate. Relative gene expression was calculated using the ΔΔCt method and normalized to the geometric mean of two housekeeping genes (Hprt1 and Actb). A complete list of TaqMan Gene Expression Assays used in this study is provided in Supplementary Table 1.

### RNA sequencing: library preparation and sequencing

For bulk RNA sequencing, total RNA was extracted from frozen punches of the nucleus accumbens (NAc), prefrontal cortex (PFC), basolateral amygdala (BLA), and ventral hippocampus (vHIP) using the RNeasy Micro Kit (Qiagen) according to the manufacturer’s instructions, including on-column DNase digestion. For viral overexpression cohorts, bilateral punches of virally infected NAc core and medial shell tissue were collected under fluorescent illumination from 1-mm coronal sections using a 14-gauge needle and frozen on dry ice. RNA concentration and integrity were assessed using an Agilent Bioanalyzer, and samples with RNA integrity number (RIN) values ≥7 were used for downstream processing.

For the female CSDS cohort, stranded total RNA libraries were prepared by Azenta/GENEWIZ using the SMARTer Stranded Total RNA Sample Prep Kit for mammalian samples. Libraries were Illumina-compatible, dual-indexed, and sequenced on an Illumina NovaSeq platform using 2 × 150 bp paired-end reads. For female and male viral overexpression cohorts, stranded total RNA libraries were prepared using the SMARTer Stranded Total RNA-Seq Kit v3 Pico Input Mammalian (TaKaRa Biotech), with ribosomal RNA depletion, according to the manufacturer’s instructions. Libraries were sequenced by Azenta/GENEWIZ on an Illumina NovaSeq platform using 2 × 150 bp paired-end reads, targeting approximately 30–40 million reads per sample. All samples within a given experiment were multiplexed and sequenced together to minimize batch effects. Raw sequencing reads were subjected to quality control using FastQC v0.11.9 (www.bioinformatics.babraham.ac.uk/projects/fastqc/). All raw sequencing reads underwent adapter trimming and were aligned to the mouse reference genome (GRCm38/mm10) using Trimmomatic v0.39 and HISAT2 v2.1.0 (daehwankimlab.github.io/hisat2/)

The top 30% most expressed annotated genes/features (highest normalized read counts average across all samples) were kept for subsequent analysis to filter out poorly expressed genes. Differential gene expression analyses were conducted in R v4.0.2 using the DESeq2 package v1.34.0^28^. Negative binomial generalized linear models were fit for each comparison of interest (e.g., RES vs CON, SUS vs CON, viral overexpression vs mCherry within sex), with experimental covariates included where appropriate. Variance-stabilizing transformation (VST) was applied to normalized count data for visualization, clustering, and network analyses. Significance cut-offs were of at least 15% expression fold change (∣log_2_(FoldChange)∣ > log_2_(1.15)) and nominal *p* < 0.05, except in the cases when genes were examined individually, where the Benjamini-Hochberg corrected *p_adj_* was used. Gene lists and corresponding statistics are available in Supplementary Table 2.

### Pattern analysis of expression trajectories

To identify recurring expression trajectories across control, susceptible and resilient mice, we performed pattern analysis using the degPatterns function from the DEGreport R package version 1.39.6 separately for each sex and brain region. For a given comparison, we first generated the union of DEGs across RES vs CON and SUS vs CON within that region. VST-normalized expression values were z-score standardized per gene across all samples. Genes were then clustered by trajectory using the degPatterns function from the DEGreport package (or equivalent clustering routine) specifying four clusters, yielding Patterns A–D. Clusters were manually annotated as resilience-specific (A,B) or stress-responsive (C,D) based on their group-wise expression profiles. Gene lists and corresponding statistics are available in Supplementary Table 2.

### Rank–rank hypergeometric overlap 2

To compare global transcriptional signatures across sex, we used rank–rank hypergeometric overlap analysis 2 (RRHO2)^19^ (<u>github.com/RRHO2/RRHO2</u>). For each region and phenotype (RES vs CON, SUS vs CON), genes were ranked by signed –log10(P) (direction given by log2FC). RRHO2 maps were generated in R using the RRHO2 package with default parameters, producing two-dimensional heatmaps representing the significance of overlapping up- or downregulated genes between female and male datasets.

### Gene ontology and pathway analyses

Gene ontology (GO) enrichment analyses were performed using Enrichr^29,30^ querying the Gene Ontology Biological Process 2023 database. Separate analyses were run for upregulated and downregulated DEGs in each pattern, module, or manipulation as indicated. Enrichr combined scores or –log10(adjusted P) values were used to rank terms; representative nonredundant GO terms were selected for display. Plots were made with ggplot2 (v3.4.2). Specific terms presented are summarized if redundancies were present.

Canonical pathway and upstream regulator analyses for viral overexpression datasets were performed using Ingenuity Pathway Analysis (IPA; Qiagen). DEG lists (typically |log2FC| ≥ log2(1.15), P < 0.05) were input, and IPA’s z-score and overlap P-value metrics were used to identify enriched pathways and predicted upstream regulators. Only endogenous molecules or chemicals were considered in the reported upstream regulator results.

### MEGENA co-expression network construction and module enrichment

Multiscale gene co-expression network analysis was performed using the MEGENA R package^20^ (github.com/songw01/MEGENA). Briefly, RNA-seq data were combined across all experimental groups within each sex (control, susceptible, and resilient) to construct sex- and region-specific co-expression networks. Variance-stabilized expression values were used to compute gene-to-gene Pearson correlation coefficients across all expressed genes passing minimum expression thresholds. Statistically significant correlations were identified using permutation-based false discovery rate estimation (10 permutations; FDR cutoff = 0.05), and planar filtered networks (PFNs) were generated using the calculate.PFN function. Multiscale clustering and module detection were then performed using do.MEGENA, with a minimum module size of 10 genes.

In the NAc, the female PFN was constructed from 28,666 expressed genes and contained 30,461 edges distributed across 85 modules, with absolute correlation coefficients ranging from approximately 0.66 to 1.00 (mean ≈ 0.80, s.d. ≈ 0.08). The male NAc PFN contained 34,247 edges and 187 modules, coefficients ranging from approximately 0.71 to 1.00 (mean ≈ 0.86, s.d. ≈ 0.07). Reflecting a greater degree of modular fragmentation; correlation coefficients were of comparable magnitude to those observed in females (full correlation summaries are provided in Supplementary Table 3). Modules containing fewer than 50 genes were excluded from downstream analyses.

Hub genes within each module were defined using a combination of node degree and node strength metrics, applying a 5% false discovery rate threshold based on permutation testing. To relate co-expression modules to stress-related phenotypes, module gene sets were tested for enrichment of differentially expressed genes from resilient versus control and susceptible versus control comparisons using Fisher’s exact test with Benjamini–Hochberg correction. Hierarchical module organization and DEG enrichment patterns were visualized using sunburst plots generated with the sunburstR package (github.com/timelyportfolio/sunburstR). Network statistics for all regions (NAc, PFC, BLA, and vHIPP), including numbers of samples, genes, edges, modules, and correlation summaries, are reported in Supplementary Table 3.

### Gene network visualization

Network layouts for selected MEGENA modules (e.g. female Module10, male Module2) were visualized in R using the igraph^31^ version 2.2.1 and ggraph version 2.2.2 packages. Edges reflected MEGENA-defined co-expression relationships, and node size was scaled by node strength or degree. Node color indicated sex specificity (female-only, male-only, or shared genes). Where indicated, nodes were additionally colored by log2FC from specific comparisons (e.g. RES vs CON, viral vs mCherry), with yellow and blue representing up- and downregulation, respectively.

### Behavioral and molecular statistics

Statistical analyses for behavioral and qPCR data were performed in R (v4.2.2) and/or GraphPad Prism (v10). For two-group comparisons, two-tailed unpaired t-tests (with Welch’s correction where appropriate) were used. For experiments involving multiple groups and/or factors (e.g. virus × stress), two-way or mixed-model ANOVAs were applied as appropriate, followed by Bonferroni or Tukey post hoc tests when main effects or interactions were significant. Normality and homogeneity of variance were assessed using Shapiro–Wilk and Levene tests, respectively. For data violating parametric assumptions, nonparametric tests (e.g. Kruskal–Wallis with Dunn’s post hoc comparisons) were used. Linear regression analyses were used to examine relationships between SI ratio and gene expression, with R² values reported. Exact tests, F or t values, degrees of freedom, and P values are reported in the figure legends or Supplementary Tables. All tests were two-tailed with significance set at P < 0.05. Data are presented as mean ± s.e.m. unless otherwise noted.

## Acknowledgements

This work was supported by the National Institute of Mental Health (R01MH129306 to E.J.N. and F31MH133297 to T.M.G.), the Hope for Depression Research Foundation (to E.J.N.), and the National Heart, Lung, and Blood Institute (T32HL007954 to T.M.G.). We thank Katherine Beach, Catherine McManus, Kyra Schmidt, Nathalia Pulido, Stephen Pirpinias, and Ezekiell Mouzon for their assistance with animal husbandry. We are grateful to Dr. Logan Brown and Dr. Boris Kantor at the Duke University Viral Vector Core for cloning and packaging AAV vectors. We also thank Clementine Blaschke and Kinneret Rosen for their contributions to laboratory operations and experimental support.

## Author contributions

Conceptualization: TMG, EP, LH, LP, EJN

Methodology: TMG, EP, LH, LP, AR, ME

Investigation: TMG, EP, LH, LP, RF, AG, CB, BK, AM, TM, TD, RD, LL, AR, ME, MR, YY, AC, OS, CN

Formal analysis: TMG, LH, AG

Visualization: TMG, LH

Funding acquisition: TMG, EJN

Supervision: EJN, EP, LH, SR, BZ, LS

Writing – original draft: TMG

Writing – review & editing: TMG, EJN

## Data availability

The RNA-seq data reported in the paper are deposited in GEO with the accession numbers GSE118317, GEO: GSE72343, & additional GEO numbers will be added before publication. Other data that support the findings of this study are available from the corresponding author upon request.

## Code availability

Scripts and code utilized in the analysis of study data are available from the corresponding

## References

1. Southwick, S. M., Charney, D. & DePierro, J. M. *Resilience: The Science of Mastering Life’s Greatest Challenges*. (Cambridge University Press, Cambridge, United Kingdom New York, NY, 2023). doi:10.1017/9781009299725.

2. Akil, H. & Nestler, E. J. The neurobiology of stress: Vulnerability, resilience, and major depression. Proc. Natl. Acad. Sci. 120, e2312662120 (2023).

3. Lorsch, Z. S. et al. Stress resilience is promoted by a Zfp189-driven transcriptional network in prefrontal cortex. Nat. Neurosci. 22, 1413–1423 (2019).

4. Bagot, R. C. et al. Circuit-wide transcriptional profiling reveals brain region-specific gene networks regulating depression susceptibility. Neuron 90, 969–983 (2016).

5. Nestler, E. J. & Russo, S. J. Neurobiological basis of stress resilience. Neuron 112, 1911–1929 (2024).

6. Navarrete, J. et al. Individual Differences in Volitional Social Self-Administration and Motivation in Male and Female Mice Following Social Stress. Biol. Psychiatry 96, 309–321 (2024).

7. Kessler, R. C., Chiu, W. T., Demler, O. & Walters, E. E. Prevalence, Severity, and Comorbidity of 12-Month DSM-IV Disorders in the National Comorbidity Survey Replication. Arch. Gen. Psychiatry 62, 617–627 (2005).

8. Perugi, G. et al. Gender-Mediated Clinical Features of Depressive Illness the Importance of Temperamental Differences. Br. J. Psychiatry 157, 835–841 (1990).

9. Mansouri, S. et al. Transcriptional dissection of symptomatic profiles across the brain of men and women with depression. Nat. Commun. 14, 6835 (2023).

10. Labonté, B. et al. Sex-Specific Transcriptional Signatures in Human Depression. Nat. Med. 23, 1102–1111 (2017).

11. Traumatic Stress Brain Research Group et al. Transcriptomic organization of the human brain in post-traumatic stress disorder. Nat. Neurosci. 24, 24–33 (2021).

12. Seney, M. L. et al. Opposite Molecular Signatures of Depression in Men and Women. Biol. Psychiatry 84, 18–27 (2018).

13. Montgomery, K. R. et al. Chemogenetic activation of CRF neurons as a model of chronic stress produces sex-specific physiological and behavioral effects. Neuropsychopharmacology 49, 443–454 (2024).

14. Cissé, Y. M. et al. Maternal preconception stress produces sex-specific effects at the maternal:fetal interface to impact offspring development and phenotypic outcomes†. Biol. Reprod. 110, 339–354 (2024).

15. Gyles, T. M., Nestler, E. J. & Parise, E. M. Advancing preclinical chronic stress models to promote therapeutic discovery for human stress disorders. Neuropsychopharmacology 49, 215–226 (2024).

16. Krishnan, V. et al. Molecular Adaptations Underlying Susceptibility and Resistance to Social Defeat in Brain Reward Regions. Cell 131, 391–404 (2007).

17. Berton, O. et al. Essential Role of BDNF in the Mesolimbic Dopamine Pathway in Social Defeat Stress. Science 311, 864–868 (2006).

18. Takahashi, A. et al. Establishment of a repeated social defeat stress model in female mice. Sci. Rep. 7, 12838 (2017).

19. Rosenblatt, J. & Stein, J. RRHO: Test overlap using the Rank-Rank Hypergeometric test. (2014).

20. Song, W.-M. & Zhang, B. Multiscale Embedded Gene Co-expression Network Analysis. PLOS Comput. Biol. 11, e1004574 (2015).

21. Wang, J. et al. A multi-omic approach implicates novel protein dysregulation in post-traumatic stress disorder. Genome Med. 17, 43 (2025).

22. Arcego, D. M. et al. Sex-specific interaction effects of Syntaxin 1A coexpression network and childhood trauma on adult depressive symptoms. eBioMedicine 123, 106062 (2026).

23. Schiweck, C. et al. Childhood trauma, suicide risk and inflammatory phenotypes of depression: insights from monocyte gene expression. Transl. Psychiatry 10, 296 (2020).

24. Islam, M. K. et al. Integrated bioinformatics and statistical approach to identify the common molecular mechanisms of obesity that are linked to the development of two psychiatric disorders: Schizophrenia and major depressive disorder. PLOS ONE 18, e0276820 (2023).

25. Song, J. & Kim, Y. Animal models for the study of depressive disorder. CNS Neurosci. Ther. 27, 633–642 (2021).

26. Hashimoto, R. et al. The Breakpoint Cluster Region Gene on Chromosome 22q11 is Associated with Bipolar Disorder. Biol. Psychiatry 57, 1097–1102 (2005).

27. Hamilton, P. J., Lim, C. J., Nestler, E. J. & Heller, E. A. Viral Expression of Epigenome Editing Tools in Rodent Brain Using Stereotaxic Surgery Techniques. in Epigenome Editing (eds Jeltsch, A. & Rots, M. G.) vol. 1767 205–214 (Springer New York, New York, NY, 2018).

28. Love, M. I., Huber, W. & Anders, S. Moderated estimation of fold change and dispersion for RNA-seq data with DESeq2. Genome Biol. 15, 550 (2014).

29. Xie, Z. et al. Gene Set Knowledge Discovery with Enrichr. Curr. Protoc. 1, e90 (2021).

30. The Gene Ontology Consortium. The Gene Ontology resource: enriching a GOld mine. Nucleic Acids Res. 49, D325–D334 (2021).

31. Csardi, G. & Nepusz, T. The igraph software package for complex network research. InterJournal (2006).

