## Supplemental Figures for "A shared transcriptional network in the nucleus accumbens supports resilience to chronic stress across sex"

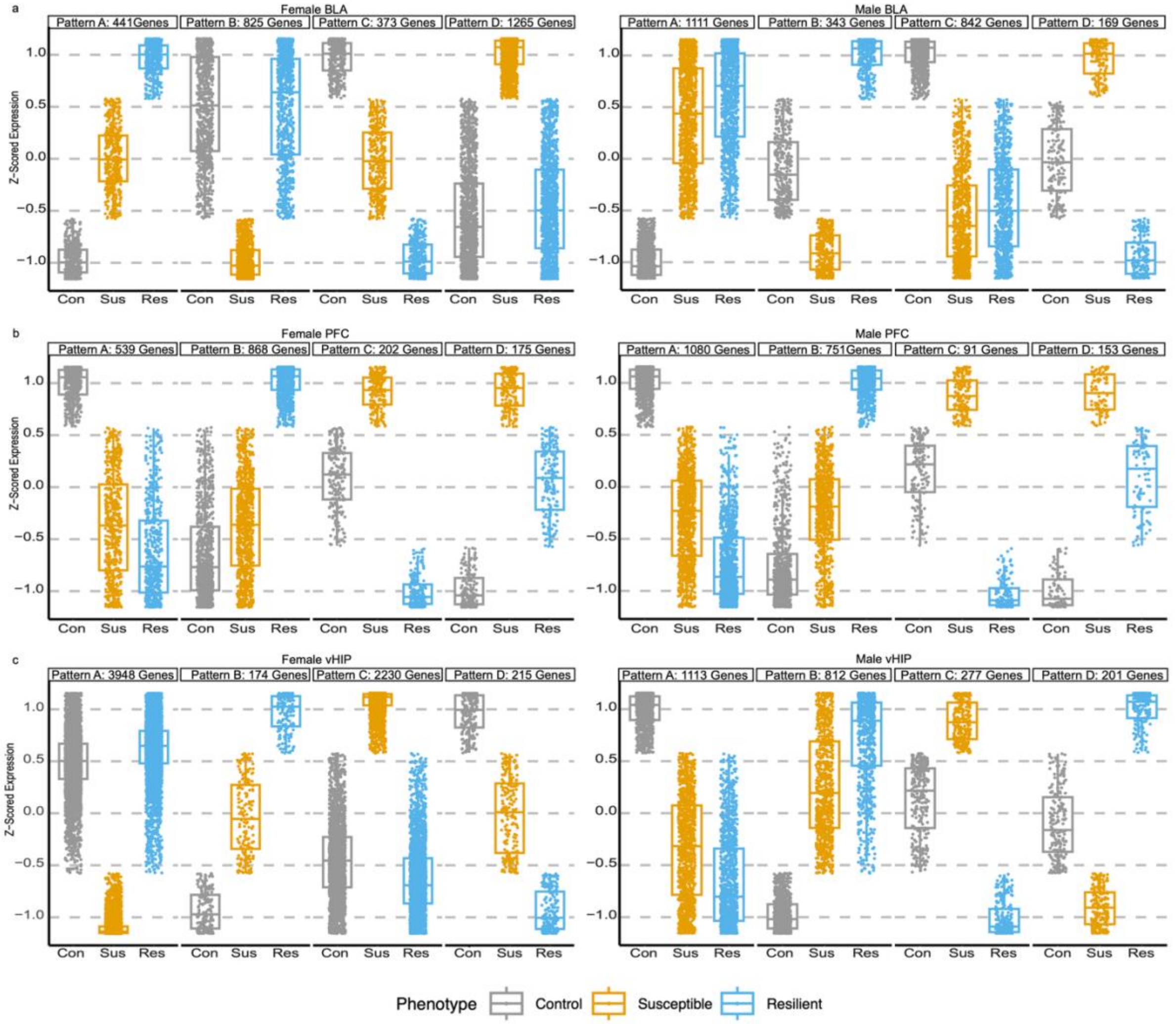


Supplementary Fig. 1. Pattern analysis of differentially expressed genes across limbic brain regions and sex.

**a**, Z-scored expression trajectories of differentially expressed genes (DEGs) in the basolateral amygdala (BLA) of female (left) and male (right) mice following chronic social defeat stress (CSDS). DEGs were grouped into four expression patterns (Patterns A–D) based on their expression profiles across control (CON), susceptible (SUS), and resilient (RES) mice. In females, Patterns A and C reflect stress-responsive expression changes, whereas Patterns B and D show susceptibility-specific regulation. In males, Patterns A and C are stress responsive, while Patterns B and D exhibit bidirectional expression changes across phenotypes. Numbers above each panel indicate the number of genes assigned to each pattern.

**b**, Z-scored expression trajectories of DEGs in the prefrontal cortex (PFC) of female (left) and male (right) mice following CSDS. In both sexes, Patterns A and D represent stress-responsive expression changes, Pattern B is enriched for resilience-specific regulation, and Pattern C shows bidirectional expression across phenotypes. Although the overall pattern structure is similar across sexes, patterns were defined independently for females and males and contain largely distinct gene sets, consistent with limited transcriptional overlap observed at the gene level. Gene counts for each pattern are shown above the panels.

**c**, Z-scored expression trajectories of DEGs in the ventral hippocampus (vHIP) of female (left) and male (right) mice following CSDS. In females, Patterns A and C are susceptibility-specific, while Patterns B and D are stress responsive. In males, Patterns A and B are stress responsive, whereas Patterns C and D display bidirectional expression changes. As in other regions, patterns were defined independently by sex, and numbers above each panel indicate the number of genes per pattern.

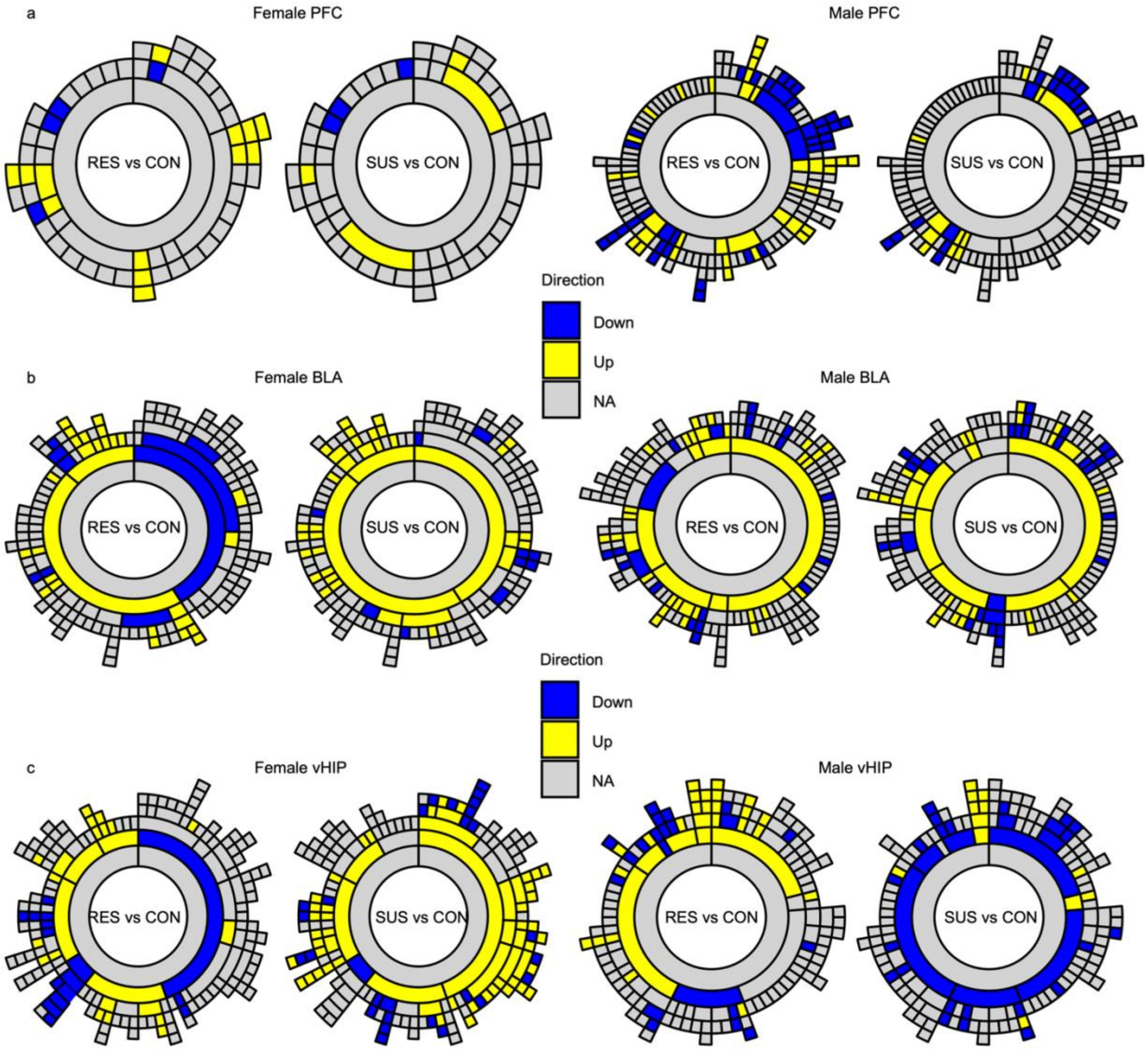


**Supplementary Fig. 2.** MEGENA module enrichment across brain regions and sex.

**a**, Sunburst plots showing enrichment of differentially expressed genes (DEGs) within prefrontal cortex (PFC) MEGENA modules for female (left) and male (right) mice. In females, overall enrichment is relatively low and distributed similarly across resilient versus control (RES vs CON) and susceptible versus control (SUS vs CON) comparisons. In males, moderate enrichment is observed, with a greater prevalence of downregulated DEGs in the RES vs CON comparison relative to SUS vs CON.

**b**, Sunburst plots showing enrichment of DEGs within basolateral amygdala (BLA) MEGENA modules for female (left) and male (right) mice. In females, strong enrichment is observed in both phenotypes, characterized by pronounced downregulation in RES vs CON and a strong signal of upregulation in SUS vs CON; a subset of modules shows shared upregulation across both phenotypes. In males, robust enrichment is evident, with broadly similar enrichment patterns across RES vs CON and SUS vs CON comparisons.

**c**, Sunburst plots showing enrichment of DEGs within ventral hippocampus (vHIP) MEGENA modules for female (left) and male (right) mice. Strong enrichment is observed across both phenotypes and sexes. In females, RES vs CON comparisons are dominated by downregulated DEGs, whereas SUS vs CON comparisons show strong upregulation, with some modules exhibiting shared upregulation across phenotypes. In males, RES vs CON is characterized by prominent enrichment of upregulated DEGs, while SUS vs CON shows enrichment largely driven by downregulated DEGs.

In all panels, inner rings represent higher-order modules and outer rings represent nested child modules. DEG directionality is indicated by color (yellow, upregulated; blue, downregulated; gray, not significant).

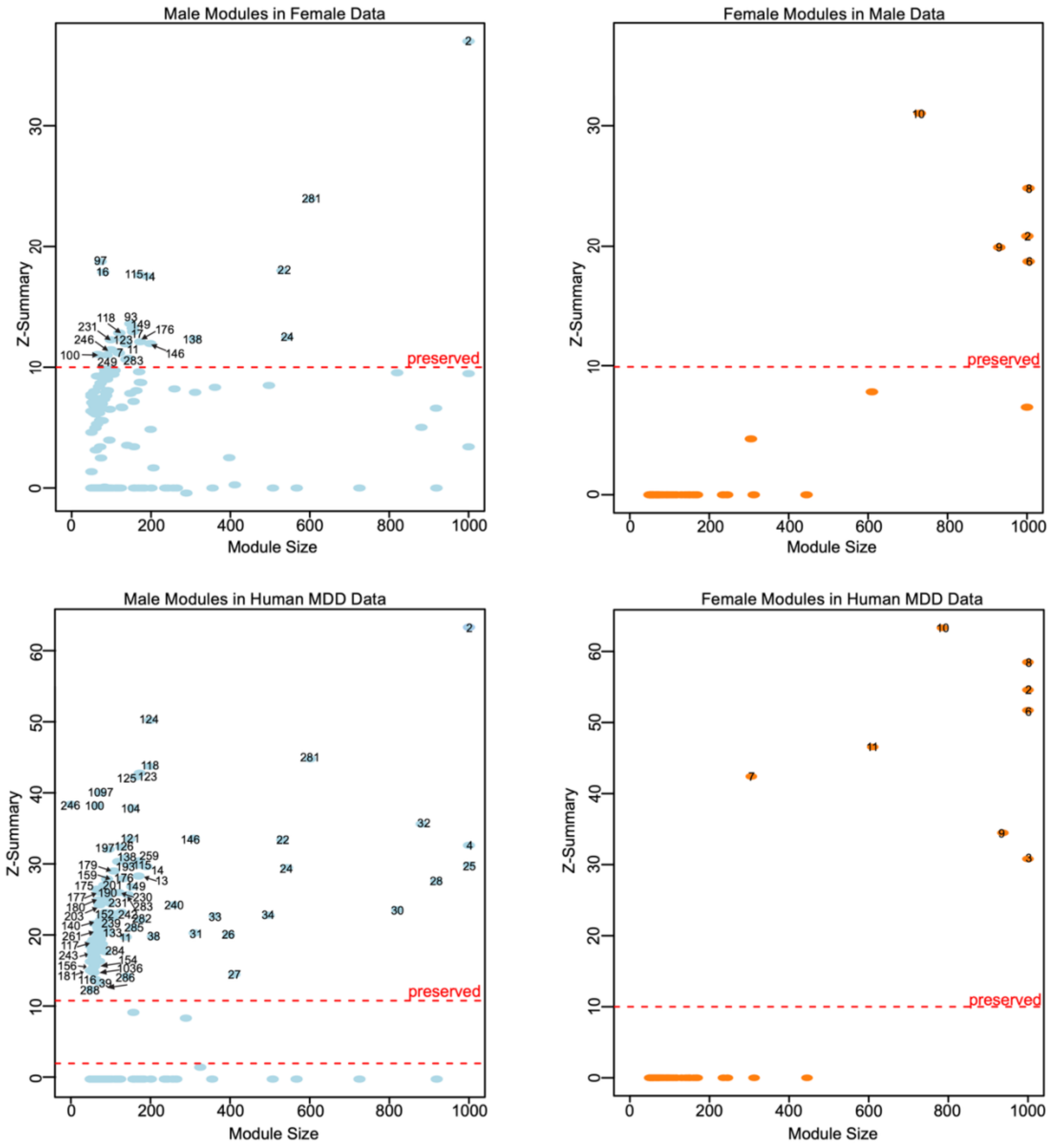


**Supplementary Fig. 3**. Module preservation analysis across sex and species.

Scatter plots show MEGENA-derived nucleus accumbens (NAc) module preservation statistics (Zsummary) plotted against module size. The dashed red line indicates the threshold for strong preservation (Zsummary ≥ 10).

**Top left**, Preservation of male mouse NAc modules in female mouse data. Using the preservation threshold, 24 of 187 male modules (12.8%) were strongly preserved in females. Although a relatively large number of male modules met the preservation criterion, most preserved modules were small, typically containing fewer than 200 genes, and together encompassed 6,734 genes.

**Top right**, Preservation of female mouse NAc modules in male mouse data. Five of 85 female modules (5.8%) were strongly preserved in males. In contrast to the male-to-female comparison, preserved female modules were consistently large, generally exceeding 600 genes in size, and together encompassed 7,245 genes.

**Bottom left**, Preservation of male mouse NAc modules in human major depressive disorder (MDD) datasets. Ninety-three of 187 male modules (49.7%) met the preservation threshold in human data. Similar to the mouse cross-sex comparison, preserved male modules tended to be smaller in size and together encompassed 13,183 mouse genes.

**Bottom right**, Preservation of female mouse NAc modules in human MDD datasets. Eight of 85 female modules (9.4%) were strongly preserved in human data. As observed in the mouse comparison, preserved female modules were among the largest female-derived modules and together encompassed 6,811 mouse genes. Preservation in human MDD datasets was assessed against pooled male and female human samples; the greater number of male samples in the human dataset may contribute to the higher apparent preservation of male mouse modules.

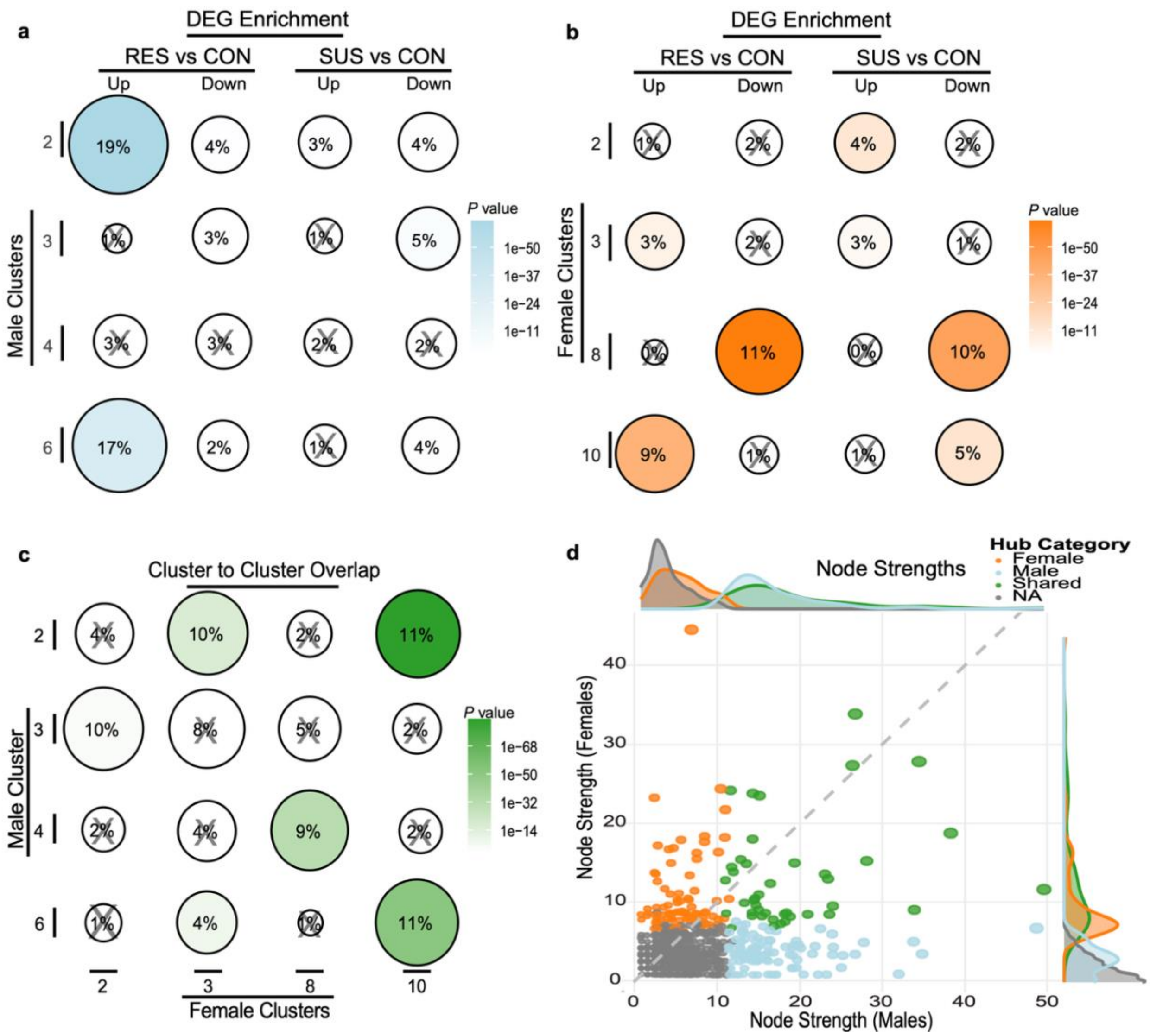


**Supplementary Fig. 4**. Module-level enrichment and cross-sex correspondence of resilience-associated MEGENA clusters.

**a**, Bubble plots showing enrichment of differentially expressed genes (DEGs) within selected male nucleus accumbens (NAc) MEGENA modules for resilient versus control (RES vs CON) and susceptible versus control (SUS vs CON) comparisons. Bubble size and percentage indicate the Jaccard index between module gene sets and DEGs, while color intensity reflects enrichment p-values. Modules marked with an "X" did not meet enrichment criteria due to non-significant p-values or odds ratios ≤1. Male modules 2 and 6 show the highest DEG enrichment.

**b**, Bubble plots showing DEG enrichment within selected female NAc MEGENA modules. Female module 8 exhibits shared downregulation across RES vs CON and SUS vs CON comparisons, whereas module 10 shows enrichment for upregulated DEGs in RES vs CON and downregulated DEGs in SUS vs CON.

**c**, Bubble plot showing cross-sex module-to-module overlap between male and female NAc MEGENA modules. Male module 2 and female module 10 show the highest cross-sex enrichment, as indicated by the largest Jaccard indices and most significant p-values.

**d**, Scatter plot showing node strength values for genes within male module 2 and female module 10. Each point represents a gene, plotted by node strength in the male network (x-axis) versus the female network (y-axis). Genes are colored by module origin (blue, male-specific; orange, female-specific; green, shared hub genes identified in both datasets). Marginal density plots illustrate the distribution of node strengths across sexes.
